# A window into lysogeny: Revealing temperate phage biology with transcriptomics

**DOI:** 10.1101/787010

**Authors:** Siân V. Owen, Rocío Canals, Nicolas Wenner, Disa L. Hammarlöf, Carsten Kröger, Jay C. D. Hinton

## Abstract

Integrated phage elements, known as prophages, are a pervasive feature of bacterial genomes. Prophages can enhance the fitness of their bacterial hosts by conferring beneficial functions, such as virulence, stress tolerance or phage resistance, which are encoded by accessory loci. Whilst the majority of phage-encoded genes are repressed during lysogeny, accessory loci are often highly expressed. However, novel prophage accessory loci are challenging to identify based on DNA sequence data alone. Here, we use bacterial RNA-seq data to examine the transcriptional landscapes of five *Salmonella* prophages. We show that transcriptomic data can be used to heuristically enrich for prophage features that are highly expressed within bacterial cells and often represent functionally-important accessory loci. Using this approach we identify a novel anti-sense RNA species in prophage BTP1, STnc6030, which mediates superinfection exclusion of phage BTP1 and immunity to closely-related phages. Bacterial transcriptomic datasets are a powerful tool to explore the molecular biology of temperate phages.

## INTRODUCTION

Temperate bacteriophages can incorporate their chromosomes into the genome of host bacteria (lysogeny), and persist as vertically replicated genomic elements known as prophages. The majority of the temperate phage genome encodes proteins dedicated to virion production and host cell lysis, functions that are toxic to bacterial cells. To exist stably as passive genomic elements, the expression of the majority of prophage genes must be repressed at the transcriptional level. Only those functions necessary to maintain and favour lysogeny are expressed. The molecular mechanisms governing prophage gene regulation are best understood in the archetypical *Escherichia coli* phage lambda, in which the synthesis of a single protein, the CI repressor, is sufficient to maintain the lysogenic state of the prophage (Ptashne, 2004). Molecular studies investigating the gene expression of phage lambda lysogens showed that four additional lambda genes are expressed from the integrated prophage genome during lysogenic growth: the *rexAB* operon, encoding a superinfection immunity system; and the *lom* and *bor* genes, which encode virulence factors (Barondess and Beckwith, 1995; Chen et al., 2005; Liu et al., 2013; Osterhout et al., 2007; Vica Pacheco et al., 1997). The expression of fitness-associated prophage-encoded genes during lysogeny represents a mutualism whereby enhancing the fitness of the bacterial host directly increases the fitness (as measured by genome replication) of the integrated prophage (Cumby et al., 2012). Such genes, known as accessory or moron loci (Juhala et al., 2000), with diverse functions such as virulence, stress resistance and phage resistance have been found in characterised bacterial prophages (Casjens and Hendrix, 2015; Cumby et al., 2012; Veses-Garcia et al., 2015). Importantly, prophage accessory loci are not required for any part of the phage life cycle, and only affect the biology of the host bacterium during lysogeny, a process known as lysogenic conversion. In *Salmonella enterica* serovar Typhimurium (*S.* Typhimurium), and many other bacterial pathogens, prophage-encoded accessory loci are important for virulence in animal models (Brüssow et al., 2004; Figueroa-Bossi and Bossi, 1999; Fortier and Sekulovic, 2013; Wahl et al., 2019).

The majority of bacterial taxa contain prophages (Roux et al., 2015; Touchon et al., 2016), and prophages are very frequently the main sources of genetic diversity between closely-related bacterial strains or pathovariants (Ashton et al., 2017; Mottawea et al., 2018). Given that more genome sequences exist for bacteria than for any other domain of life, and bacteria frequently harbour multiple prophages, it has been argued that temperate phage may be the most deeply sequenced organisms on the planet (Owen et al., 2018). Prophages therefore represent a vast and largely unexplored reservoir of functionally important genes. However, identifying prophage accessory loci in bacterial genome sequences, particularly in bacterial taxa in which prophages have not been well characterised, is challenging. The functional gene annotation of prophage regions most frequently consists of hypothetical proteins, and it is difficult to distinguish hypothetical proteins that represent novel accessory genes from those that represent divergent genes involved in the phage lifecycle based on DNA sequence data alone.

Here we use transcriptomic data to show that prophage accessory loci typically have unique transcriptional signatures compared with phage lytic genes, an observation which can be exploited to heuristically enrich for novel prophage accessory features and genes involved in the maintenance of lysogeny. Using this approach we discovered that the uncharacterised STnc6030 anti-sense RNA (asRNA) of *S.* Typhimurium prophage BTP1 functions as a novel superinfection exclusion factor. Bacterial transcriptomic data mapped to prophage regions are a powerful tool to discover novel temperate phage biology simply by identifying the subset of prophage genetic features that are highly expressed in lysogenic cells.

## METHODS

### Transcriptomic analysis of the prophage regions of *S.* Typhimurium D23580

Prophage gene expression during lysogeny was investigated using data from three previous studies (Canals et al., 2019; Hammarlöf et al., 2018; Kröger et al., 2013). A description of the experimental conditions associated with the RNA-seq data used in this analysis is given in Supplementary Table 1. Transcripts per million (TPM) values were obtained from Canals et al. and represent a normalised expression value per gene and condition (Canals et al., 2019; Wagner et al., 2013, 2012). As previously described, a TPM cut-off score of 10 was used to define gene expression, so that only genes with a TPM value of >10 were considered to be expressed (Kröger et al., 2013). Heat maps showing absolute expression were obtained using TPM values and conditional formatting in Microsoft Excel. Prophage transcriptome maps were generated by visualisation of sequence reads using the Integrated Genome Browser (IGB) (Nicol et al., 2009). For display in IGB, the read depth was adjusted in relation to the cDNA library with the lowest number of reads (Skinner et al., 2009). For identification of prophage regulatory or accessory genes, a TPM cut-off score of 100 was used to define high expression, so that only prophage genes with a TPM value of >100 were considered to be highly expressed and therefore putatively involved in prophage lysogeny regulation or accessory functions.

### Plasmid construction

All oligonucleotide (primer) sequences used in this study are listed in Supplementary Table 2, and bacterial strains, phages and plasmids are given in Supplementary Table 3. The plasmid to overexpress the STnc6030 asRNA (pP_L_-STnc6030) was constructed using the overlap-extension PCR cloning method previously described (Bryksin and Matsumura, 2010).

The pJV300 (pP_L_) plasmid (Sittka et al., 2007) was initially modified to encode gentamicin resistance (pP_L_-Gm) by overlap-extension PCR cloning. For that, chimeric primers NW_88 and NW_89, containing pJV300 plasmid sequence at the 5′ end and insert sequence at the 3′ end were first used to PCR-amplify the *aacC1* gentamicin resistance locus from the pME4510 plasmid (Rist and Kertesz, 1998). 30 ng of the template plasmid were mixed with 150-300 ng of the insert and Q5 buffer, dNTPs, Q5 DNA polymerase (New England Biolabs) and water were added to a final volume of 50 μl. PCR reactions were carried out as follows: 98°C, 30 sec; 25x (98°C, 10 sec; 55°C, 30 sec, 72°C, 3 min); 72°C, 5 min. The original plasmid template was then digested using the restriction enzyme DpnI in Cutsmart buffer (1X) (New England Biolabs), according to the manufacturer’s instructions, and the overlap-extension PCR products were transformed into chemically competent *E. coli* TOP10 cells (Green and Rogers, 2013). Cells harbouring the new pP_L_-Gm (pJV300 gentamicin resistant derivative) plasmid were selected by plating on LB agar containing gentamicin (20 µg/ml). Overlap-extension PCR cloning was subsequently used to insert a sequence of interest into the pP_L_-Gm plasmid downstream of the P_LlacO-1_ constitutive promoter. The same procedure previously described was followed, except that the chimeric primers used to amplify the insert were NW_295 and NW_296, and D23580 genomic DNA was used as a template. These primers targeted the region of the pP_L_-Gm plasmid between the P_LlacO-1_ promoter and the *rrnB* transcriptional terminator. To confirm that the plasmid carried the correct construction after transformation, primers external to the insertion site were used to Sanger sequence the inserted fragment (GATC).

### Extraction of total bacterial RNA

Bacterial RNA was extracted as previously described (Kröger et al., 2013, 2012). Four or 5 OD_600_ units were removed from bacterial cultures, and cellular transcription was stopped using 0.4X culture volume of a 5% phenol (pH 4.3) 95% ethanol “stop” solution (Sigma P4557 and E7023, respectively). Cells were stabilised on ice in stop solution for at least 30 minutes before being harvested at 7,000 x *g* for 10 min at 4°C. At this point, pellets were either stored at -80°C, or RNA was immediately extracted.

To isolate RNA, pellets were resuspended in 1 ml of TRIzol^TM^ Reagent (Invitrogen). Four hundred μL of chloroform were added and the samples were immediately and thoroughly mixed by inversion. Samples were moved to a Phase-lock tube (5 Prime) and the aqueous and organic phases were separated by centrifugation at 13,000 rpm for 15 minutes at room temperature in a table top centrifuge. The aqueous phase was transferred into a new 1.5 ml tube and the RNA was precipitated using isopropanol for 30 minutes at room temperature, followed by centrifugation at 21,000 x *g* for 30 minutes at room temperature. The RNA pellet was rinsed with 70% ethanol followed by centrifugation at 21,000 x *g* rpm for 10 minutes at room temperature. The RNA pellet was air-dried for 15 minutes and resuspended in DEPC-treated water at 65°C with shaking at 900 rpm on a Thriller thermoshaker (Peqlab) for 5 minutes with occasional vortexing. RNA was kept on ice whenever possible and was stored at -80°C. RNA concentration was measured using a nanodrop ND-1000 spectrophotometer, and RNA quality was inspected visually using a 2100 Bioanalyser (Agilent).

### Detection of STnc6030 by Northern Blot

Following extraction, total RNA was separated based on size by electrophoresis through a denaturing 20 mM guanidine thiocyanate 1.5% agarose gel in TBE 1 X. Generally 1-10 μg of RNA were mixed with an equal volume of 2X Urea-Blue denaturing buffer (0.025% xylene cyanol, 0.025% bromophenol blue and 50% urea) and samples were heat-denatured at 90°C for 5 minutes and chilled on ice before loading. 4 μl of Low Range ssRNA ladder (NEB) or 5 μl RNA molecular weight marker I DIG-labelled (Roche) were treated in the same way as the RNA samples to allow the detected transcript length to be estimated. Samples were run in 1X TBE buffer at a constant voltage of 120 V for denaturing polyacrylamide gels, or 80 V (at 4°C) for denaturing agarose gels.

Separated RNA was transferred from polyacrylamide gels to positively charged nylon membranes (Roche, cat. 11 209 272 001) using the Trans-Blot SD Semi-Dry Electrophoretic Transfer Cell (BioRad) at a constant amplitude of 125 mA for 30 minutes at 4°C. For denaturing agarose gels, separated RNA was transferred to positively charged nylon membranes using overnight capillary transfer in 20X saline-sodium citrate (SSC) buffer as described in the DIG Northern Starter Kit manual (Roche, Cat. 12039672910).

RNA was UV-crosslinked to the membranes in a CL-1000 UV-crosslinker (UVP) set to 3600 (360,000 μJ/cm^2^). The membrane was equilibrated in hybridisation buffer for 1 hour at 68°C in pre-warmed DIG Easy Hyb solution (Roche) in a rotating hybridisation oven. 5 μl (approximately 1.25 μg) of riboprobe were heat denatured in 5 ml of DIG Easy Hyb solution at 68°C for 30 minutes and added to the membrane for hybridisation overnight at 68°C in the rotating hybridisation oven. The oligonucleotide sequence used to generate the DIG labelled riboprobe are given in Supplementary Table 2. Membrane washing, blocking and transcript detection steps were carried out as described in the Roche manual. An ImageQuant LAS4000 Imager was used for blot detection.

### Phage enumeration and plaquing

Phage enumeration and plaque isolations were carried out using the double layer agar technique. To count spontaneously induced phages in overnight cultures, 1 ml of overnight culture was passed through a 0.22 μm syringe filter. For enumeration, phage lysates or culture supernatants serial 10-fold dilutions were made in sterile LB broth. For overnight culture supernatant, dilution up to 10^-7^ was sufficient, whereas for phage lysates, higher dilutions were required (typically up to 10^-10^). 4 ml of 0.4% LB agar were seeded with 100 μl of an overnight culture of the required indicator strain (approximately 10^8^ CFU) and, once solidified, phage dilutions were applied to the agar in 10 μl drops. After drying for 30 mins, plates were incubated overnight at 37°C. Phage concentrations were calculated as plaque forming units (PFU) per ml of lysate or culture supernatant (PFU/ml).

### Isolation and sequencing of STnc6030 escape phage

One hundred ul of 10-fold dilutions of high titre BTP1 stock (10^11^ PFU/ml) were mixed with 100 µl of D23580 ΔBTP1 pP_L_-STnc6030, added to 3 ml of molten LB (Lennox) 0.3% agar and poured onto LB plates. The plates were incubated right-side up overnight at 37°C for 16 hours. The frequency of occurrence of spontaneous STnc6030 escape mutants was determined as the PFU/ml of escape mutants divided by the input titre of BTP1 phage. Escape phage plaques were picked and replicated on D23580 ΔBTP1 pP_L_-STnc6030. To ensure purity, a nested PCR strategy was used to ensure amplification of the STnc6030 region from the escape phages without amplification of WT STnc6030 from contaminating pP_L_-STnc6030 plasmid DNA. The larger STnc6030 region was first amplified using oligonucleotides NW_296 and NW_298, and this amplicon was used for a more targeted amplification of the STnc6030 region using oligonucleotides NW_296 and NW_295, yielding a 787 bp amplicon that was Sanger sequenced by GATC Biotech AG (Germany). All amplicons were sequenced with oligonucleotides NW_296 and NW_295 so that the entire length of the STnc6030 region could be resolved. SNPs were detected by alignment of the Sanger sequencing data with the STnc6030 sequence from WT *S.* Typhimurium D23580 using SnapGene software (from GSL Biotech; available at snapgene.com).

## RESULTS

The *S.* Typhimurium strain D23580 is a representative of the ST313 lineage 2 which is currently responsible for the epidemic of invasive non-typhoidal Salmonellosis in Africa (Kingsley et al., 2009). RNA-seq data from two previous studies (Canals et al., 2019; Hammarlöf et al., 2018) were used to interrogate the transcriptomes of the five prophages of *S.* Typhimurium D23580 (Owen et al., 2017). Further RNA-seq data for *S.* Typhimurium strain 4/74 from Kröger et al. allowed investigation of the conservation of prophage transcriptional landscapes in independent strain backgrounds (Kröger et al., 2013). Prophages Gifsy-2, ST64B and Gifsy-1 are present in both the 4/74 and D23580 strains, unlike BTP1 and BTP5, which are exclusive to strain D23580.

Our published RNA-seq experiments generated transcriptomes of bacterial cell populations grown in 17 different *in vitro* conditions (Supplementary Table 1) designed to mimic the environments that *Salmonella* experiences during infection of a mammalian host (Canals et al., 2019; Kröger et al., 2013). A differential RNA-seq (dRNA-seq) approach was used to identify transcriptional start sites (TSS) (Sharma et al., 2010) for a subset of the *in vitro* conditions, and these were included in the analysis of the prophage transcriptomes. TSS data are particularly important for prophage regions, as they indicate the position and activity of promoters required to understand the regulatory architecture of the prophages.

We previously showed that, of the five prophages in *S.* Typhimurium D23580, only prophage BTP1 produced infectious viruses (Owen et al., 2017). Prophages Gifsy-2, ST64B and Gifsy-1 all contain mutations that either prevent prophage induction (Gifsy-1), or preclude assembly of infectious viruses (ST64B and Gifsy-2). Infectious BTP5 viruses could not be detected, and therefore it is not known if the BTP5 prophage is functional. First, we used the RNA-seq data to reveal the transcriptional landscapes of these five distinct *Salmonella* prophages during lysogeny.

### Prophage transcriptional landscapes

#### Prophage BTP1

The transcriptome map (Figure 1, Supplementary Figure 1) of BTP1 in a lysogenic state showed relatively little transcription, interspersed with a small number of highly transcribed regions. To quantitatively assess prophage gene expression, we used previously established transcript per million (TPM) expression values (Canals et al., 2019). Genes with TPM values ≤10 were considered not to be expressed. Five genes (excluding the two transfer RNA (tRNA) genes located in the centre of the prophage) showed particularly high expression (TPM >100; Supplementary Table 4); *bstA* (STMMW_03531, formerly known as *ST313-td*), *cI^BTP1^*(*STMMW_03541*), *pid* (*STMMW_03751*), *gtrC^BTP1^*(*STMMW_03911*) and *gtrA^BTP1^* (*STMMW_03921*). The *cI^BTP1^* and the *bstA* genes represent an operon transcribed from a single TSS, and RNA-seq reads mapped across the 88 bp intergenic region. The *cI^BTP1^*-*bstA* operon was highly expressed in most conditions, consistent with the CI^BTP1^ protein functioning as the prophage repressor. The *pid* gene, linked to maintenance of the pseudolysogenic state in phage P22 (Cenens et al., 2013b, 2013a) also showed a high level of transcription in the majority of conditions. Lastly, a two-gene operon encoding *gtrA^BTP1^* and *gtrC^BTP1^* showed high expression (TPM >100, Supplementary Figure 1). The *gtrA^BTP1^* gene encodes a putative bactoprenol-linked glucosyltranslocase (also known as ‘flippase’), and *gtrC^BTP1^* encodes a putative acetyltransferase that mediates the addition of an acetyl group to the rhamnose subunit of the O-antigen (Kintz et al., 2015). The *gtr* operon is commonly found in P22-like phages and modifies the O-antigen component of the lipopolysaccharide to inhibit superinfection of the lysogeny (Davies et al., 2013).

**Figure 1.**
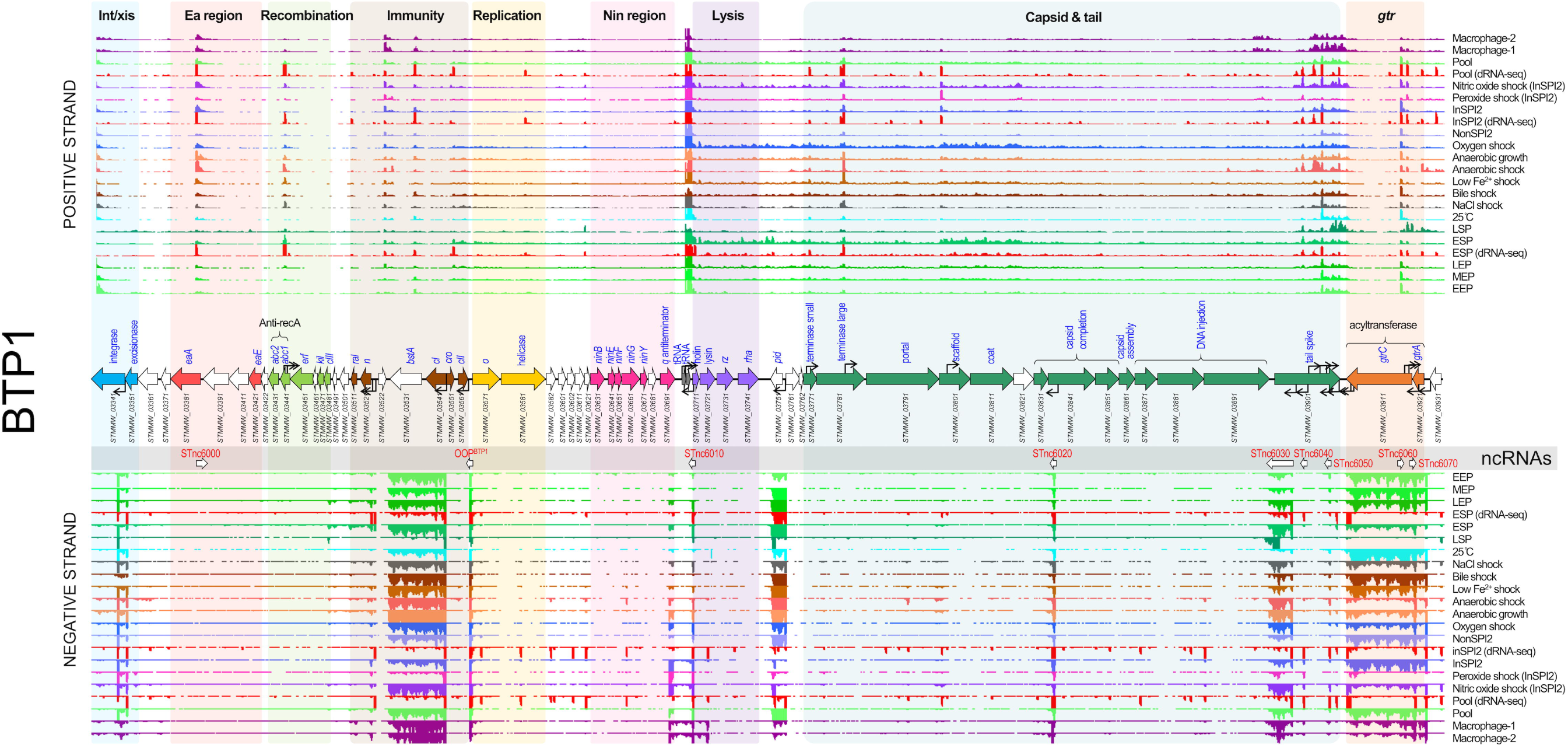
The Transcriptomic landscape of the BTP1 prophage of *S.* Typhimurium D23580 across 22 different RNA-seq experiments. RNA-seq and dRNA-seq data from Canals et al. (2019) and Hammarlöf et al. (2018). Each coloured horizontal track represents a different RNA-seq condition (Supplementary Table 1), and upper panel shows sequence reads mapped to the positive strand and lower panel, negative strand. The dRNA-seq data are shown in red, and were used to identify the TSS, which are indicated by curved black arrows on the annotation track. Annotated phage genes are grouped into functional clusters.

The structural (capsid and tail) and lysis genes (late lytic genes) showed low expression levels (TPM 10-20) in the majority of conditions tested. We previously showed that the BTP1 prophage exhibits an extremely high level of spontaneous induction (0.2%) within the cellular population (Owen et al., 2017). Because the transcriptome data generated by RNA-seq represents the average gene expression across an entire bacterial population, we propose that the apparent low-level expression of lytic genes (which are involved in cell lysis or components of the capsid and tail) reflects extremely high level transcription of lytic genes within the fraction of the population in which the BTP1 prophage is spontaneously induced. The prophage lytic genes appear to be transcribed in a large polycistronic operon that begins upstream of *STMMW_03711*, directly after the two central tRNA genes. Operons may possess more than one promoter, producing transcripts of different lengths (Kröger et al., 2012). The BTP1 prophage capsid gene cluster contained four promoters on the coding strand. The BTP1 tail spike gene showed a different transcriptional pattern to the rest of the lytic gene operon and had several secondary promoters on both the coding and non-coding strand (Figure 1).

Nine candidate noncoding RNAs (ncRNAs) were annotated in the transcriptome of BTP1 (Figure 1). One of these, designated *OOP^BTP1^*, occupies the same position as the *OOP* ncRNA of phage lambda, which is antisense to and overlapping the 3’ end of the *cII* gene (Krinke and Wulff, 1987). In phage lambda, *OOP* is thought to regulate the expression of the *cII* gene, modulating the switch between lysogenic and lytic infection (Krinke et al., 1991). The remaining nine putative ncRNAs have not been identified in other lambdoid phages and prophages, and therefore their biological function is unknown. One of the identified ncRNAs, STnc6030, was notable due to its unusually large size (>700 nt), its position antisense to phage structural genes and its high-level of expression (average TPM = 90).

#### Prophage Gifsy-2

Unlike the BTP1 prophage transcriptome, which showed expression of the lytic genes in most conditions tested, the lytic genes of Gifsy-2 were not expressed (TPM ≤10) (Figure 2). The highly expressed genes (TPM >100) of Gifsy-2 were restricted to known prophage accessory genes, such as the virulence-associated genes *sodCI* (*STMMW_10551*), *gtgE* (*STMMW_10681*) and *sseI* (*STMMW_10631*). The *sseI* gene is a known pseudogene in strain D23580 generated by the insertion of a transposase (Carden et al., 2017). Sequence redundancy of the inserted transposase gene (*STMMW_10641*) within the D23580 genome prevented the unique mapping of sequence reads, resulting in a blank region in the transcriptomic map (Figure 2).

**Figure 2.**
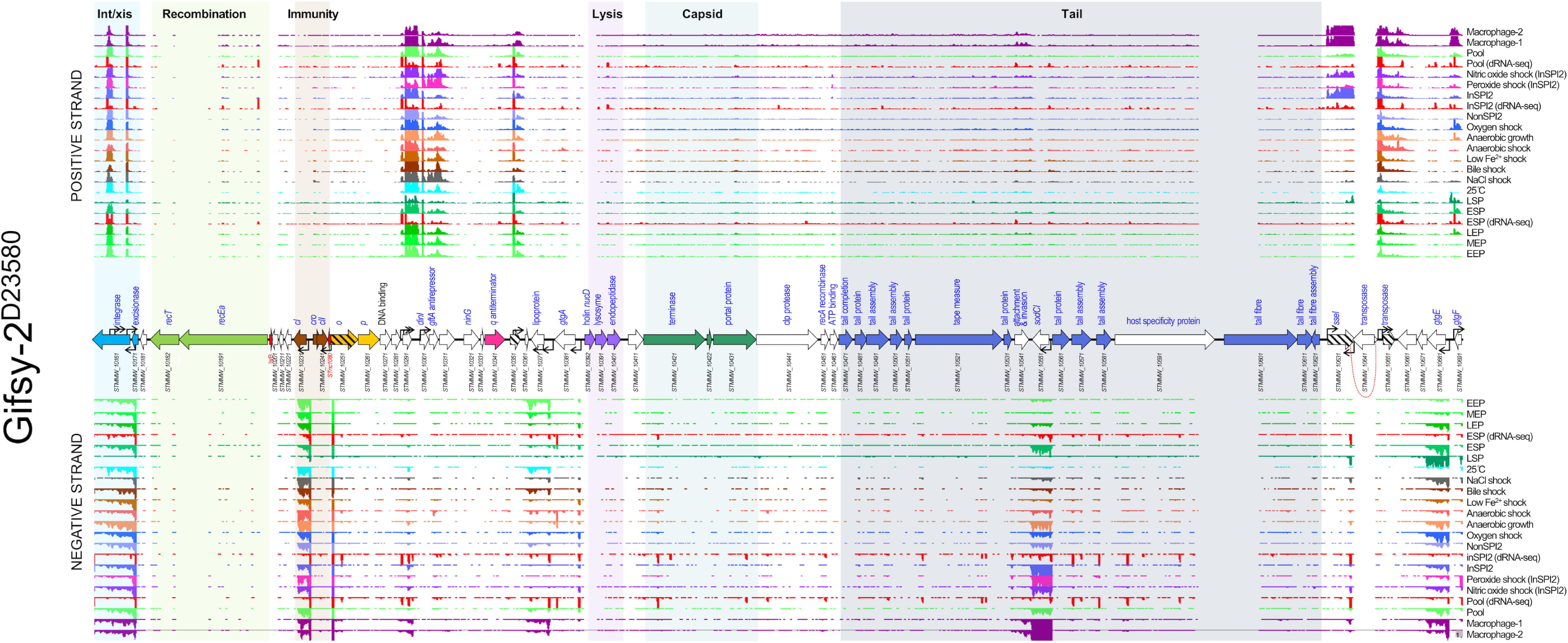
The Transcriptomic landscape of the Gifsy-2 prophage of *S.* Typhimurium D23580 across 22 different RNA-seq experiments. RNA-seq and dRNA-seq data from Canals et al. (2019) and Hammarlöf et al. (2018). Each coloured horizontal track represents a different RNA-seq condition (Supplementary Table 1), and upper panel shows sequence reads mapped to the positive strand and lower panel, negative strand. The dRNA-seq data are shown in red, and were used to identify the TSS, which are indicated by curved black arrows on the annotation track. Annotated phage genes are grouped into functional clusters. Striped arrows indicate pseudogenes and dotted red lines indicate where open reading frames have been disrupted. Due to genomic redundancy some RNA-seq reads could not be mapped uniquely to the chromosome, these reads were ignored, and so transcriptomic signal is absent from parts of the prophage (e.g. the Gifsy-2 transposase *STMMW_10641)*.

Consistent with reports that Gifsy-2 is a lambda-like prophage (Lemire et al., 2011), the Gifsy-2 homolog of the lambda CI repressor, *STMMW_10231*, was expressed (TPM >10) in all tested conditions. However, the induction behaviour of Gifsy prophages is distinct from lambda prophages. Gifsy prophages encode LexA-repressed antirepressor proteins, which, upon activation of the SOS response, inactivate the repressor protein (Lemire et al., 2011). This is opposed to the inactivation of the repressor protein by activated RecA protein in lambda prophages. We observed transcription of the Gifsy-2 antirepressor gene, *gftA*, in most of our experimental conditions (Figure 2), suggesting that additional regulatory mechanisms may be involved in the induction in the Gifsy-2 prophage.

To examine the conservation of gene expression patterns between prophages in different host backgrounds, the level of transcription of Gifsy-2 genes in *S.* Typhimurium strain D23580 (belonging to sequence type 313) was compared to the level of transcription in sequence type (ST)19 strain 4/74 (Kröger et al., 2013) (Supplementary Figure 2). The most obvious difference between the expression patterns is that the Gifsy-2 ncRNA *IsrB-1* appeared to be expressed in strain 4/74 but not in D23580. However, previous investigation showed that *IsrB-1* is duplicated in the D23850 genome (being present on both Gifsy-2^D23580^ and Gifsy-1^D23580^ prophages) (Canals et al., 2019), meaning that *IsrB-1* RNA-seq reads could not be uniquely mapped to the D23580 genome. Therefore, the absence of *IsrB-1* expression in D23580 is an artefact. Aside from this discrepancy, a remarkable consistency in the gene expression of the Gifsy-2 prophage in strains D23580 and 4/74 was observed, showing that prophage gene expression landscapes are independent of host background and highly reproducible between two phylogenetically-distinct *Salmonella* strains.

#### Prophage ST64B

The ST64B prophage transcriptome (Figure 3) only showed significant lytic-gene transcription in two of the RNA-seq conditions tested: peroxide shock and nitric oxide shock. Because hydrogen peroxide and nitric oxide cause oxidative stress and DNA damage, transcription of the lytic genes in these conditions likely represents induction of the defective ST64B prophage (the mutation in the tail assembly protein that inactivates function of the ST64B prophage is unlikely to prevent prophage induction). The lack of evidence for increased induction of the remaining D23580 prophages in these two conditions suggests that the ST64B prophage in D23580 is specifically sensitive to peroxide and nitric oxide stresses, which has not been previously observed.

**Figure 3.**
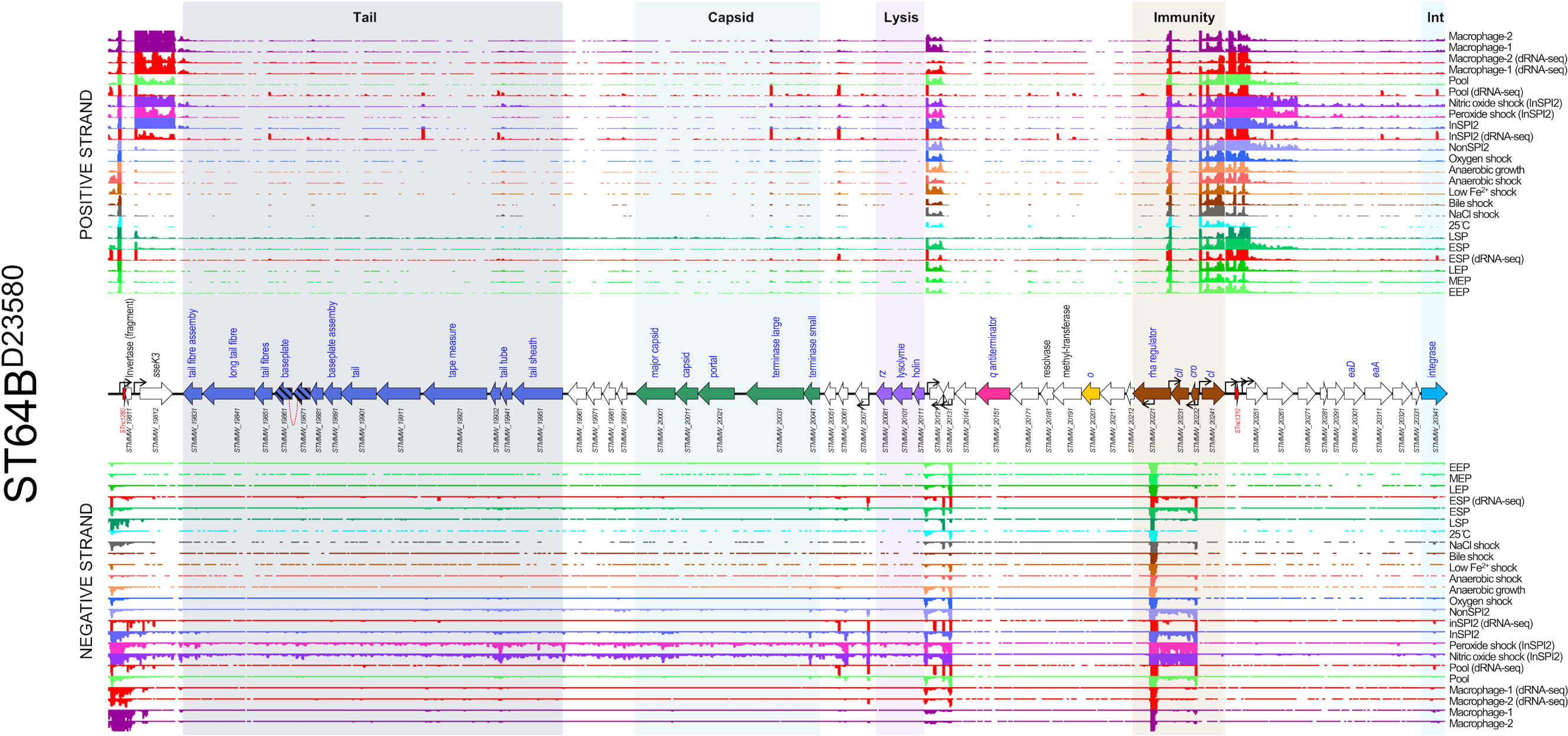
The Transcriptomic landscape of the ST64B prophage of *S.* Typhimurium D23580 across 22 different RNA-seq experiments. RNA-seq and dRNA-seq data from Canals et al. (2019) and Hammarlöf et al. (2018). Each coloured horizontal track represents a different RNA-seq condition (Supplementary Table 1), and upper panel shows sequence reads mapped to the positive strand and lower panel, negative strand. The dRNA-seq data are shown in red, and were used to identify the TSS, which are indicated by curved black arrows on the annotation track. Annotated phage genes are grouped into functional clusters.

ST64B contains an accessory gene that encodes the secreted effector protein SseK3. The *sseK3* gene was only expressed in conditions linked to infection inside mammalian macrophages, such as InSPI2 (a medium designed to mimic the intra-macrophage environment), peroxide shock, nitric oxide shock and the intra-macrophage environment. A similar expression profile was observed for genes that encode other SPI2-translocated effector proteins in *S*. Typhimurium strain 4/74 (Srikumar et al., 2015).

The patterns of gene expression of ST64B^D23580^ and ST64B^4/74^ showed many similarities (Supplementary Figure 3), though in 4/74 the prophage did not show lytic gene expression in the peroxide shock condition, suggesting that the prophage could exhibit distinct induction behaviour in strain 4/74. We speculate this difference could be explained by minor variations in the gene content between ST64B^D23580^ and ST64B^4/74^ (gene names displayed in red in Supplementary Figure 3 are unique to ST64B^D23580^). However, previous studies have shown that induction of ST64B does not produce infectious phage particles due to a frameshift mutation in the tail assembly gene (*STMMW_19861-STMMW_19871*) (Figueroa-Bossi and Bossi, 2004; Owen et al., 2017). Whilst the ST64B prophage in strain D23580 is not capable of forming infectious virions, induction of the prophage is still likely to result in cell lysis. This finding is relevant for other strains of *S.* Typhimurium which may harbour functional versions of the ST64B prophage.

#### Prophage Gifsy-1

Like the Gifsy-2^D23580^ prophage, Gifsy-1^D23580^ showed no evidence of lytic-gene transcription in any of the 17 environmental conditions examined (Figure 4). This is consistent with the very low level of spontaneous induction that was observed for the resuscitated Gifsy-1 prophage in our previous study (Owen et al., 2017). Gifsy-1^D23580^ expressed a number of ncRNAs which were highly transcribed in all conditions tested, including STnc1380, STnc2080, IsrJ, STnc1160 and IsrK (Padalon-Brauch et al., 2008). The virulence-associated genes *gogB* (*STMMW_26001*), *steE* (*STMMW_26011*, previously known as *pagJ* and *sarA*) and *pagK* (*STMMW_26041*) were only expressed in intra-macrophage infection-related conditions. In contrast, the virulence-associated gene *gipA* (*STMMW_26191*) was transcribed in all conditions tested, despite being previously reported to be specifically induced during colonisation of the small intestine (Stanley et al., 2000). Another virulence-associated gene, *gtgA* (*STMMW_26331*) showed very little transcription in any of the conditions studied. The functionally uncharacterised gene *STMMW_26411* was specifically induced in the anaerobic shock and anaerobic growth conditions, suggesting that this gene could play a role when the bacterium experiences absence of oxygen.

**Figure 4.**
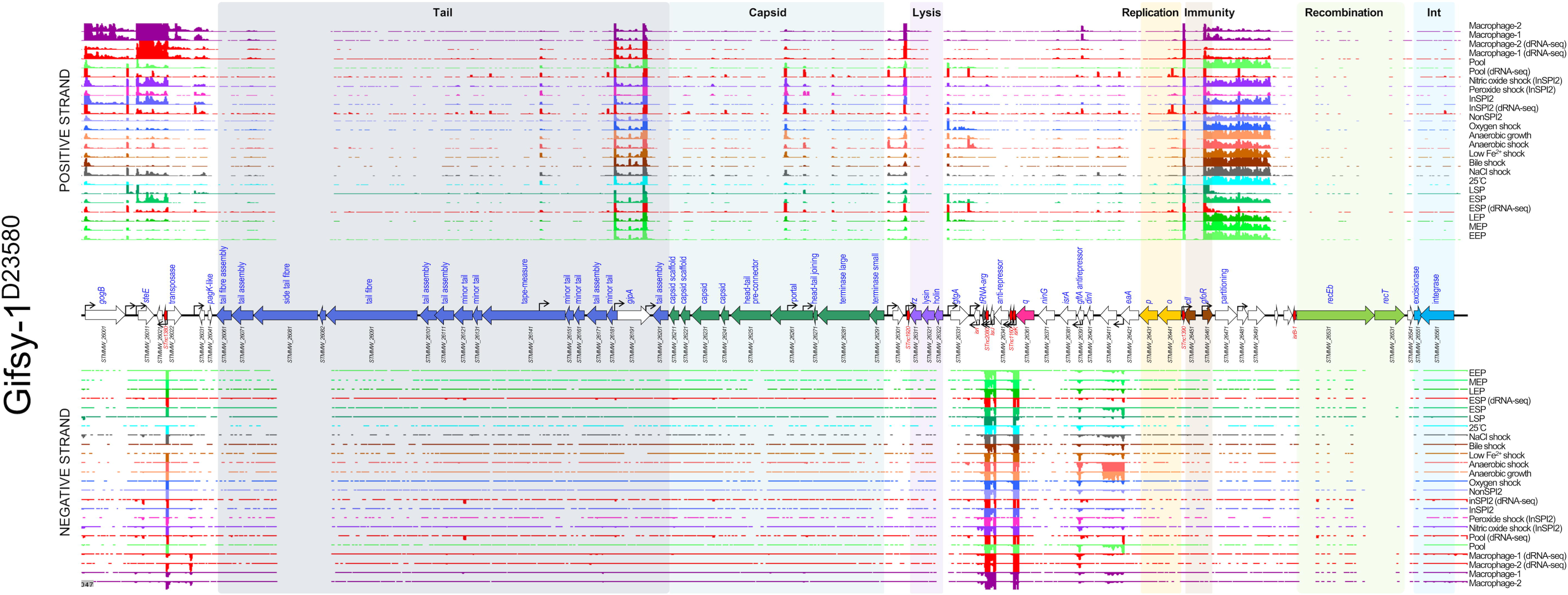
The Transcriptomic landscape of the Gifsy-1 prophage of *S.* Typhimurium D23580 across 22 different RNA-seq experiments. RNA-seq and dRNA-seq data from Canals et al. (2019) and Hammarlöf et al. (2018). Each coloured horizontal track represents a different RNA-seq condition (Supplementary Table 1), and upper panel shows sequence reads mapped to the positive strand and lower panel, negative strand. The dRNA-seq data are shown in red, and were used to identify the TSS, which are indicated by curved black arrows on the annotation track. Annotated phage genes are grouped into functional clusters.

The Gifsy-1 prophage is not identical between strains D23580 and 4/74 and shows considerable difference in gene content particularly at the 3’ terminal end. Therefore, comparison of the Gifsy-1 gene expression profiles between the two strains is difficult to interpret (Supplementary Figure 4). However, the expression patterns of orthologous genes shared by the two prophages were similar across the conditions.

#### Prophage BTP5

The BTP5 prophage showed the least transcriptional activity of all the D23580 prophages (Figure 5, Supplementary Figure 5). However, the expression pattern of BTP5 genes does not provide insight into the biology of the prophage. The most highly expressed transcript in the prophage belonged to the *tum* gene (*STMMW_32041*), a homolog of the Tum antirepressor of coliphage 186 (Shearwin et al., 1998). The antirepressor was expressed at high level particularly in the nitric oxide shock condition, raising the possibility that nitric oxide could stimulate induction of BTP5. We previously showed that infectious BTP5 phages could not be detected after induction with mitomycin C from strain D23580 (Owen et al., 2017) but the apparent activation of the antirepressor in response to hydrogen peroxide stress could suggest that the BTP5 prophage shows a specific induction behaviour.

**Figure 5.**
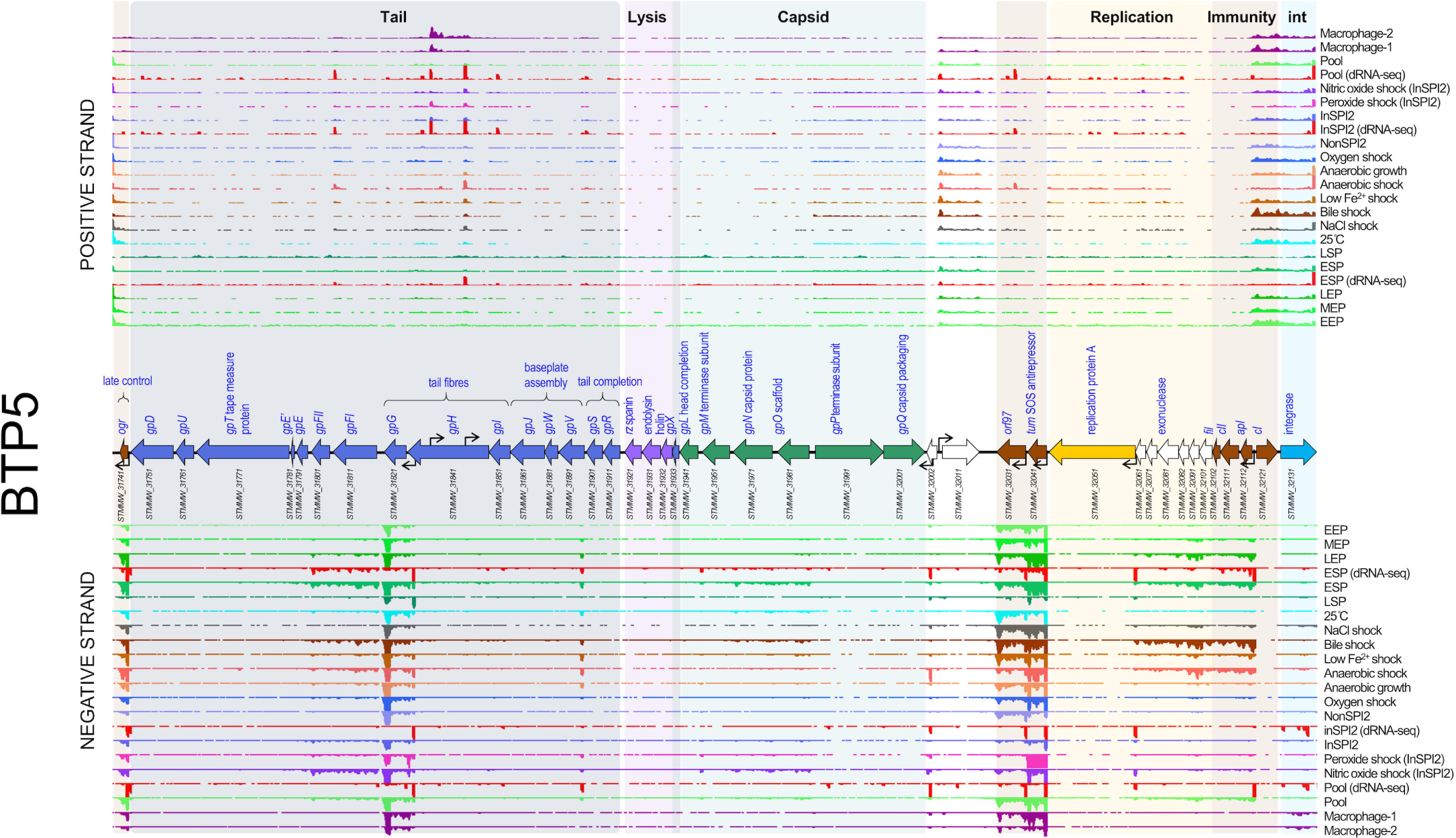
The Transcriptomic landscape of the BTP5 prophage of *S.* Typhimurium D23580 across 22 different RNA-seq experiments. RNA-seq and dRNA-seq data from Canals et al. (2019) and Hammarlöf et al. (2018). Each coloured horizontal track represents a different RNA-seq condition (Supplementary Table 1), and upper panel shows sequence reads mapped to the positive strand and lower panel, negative strand. The dRNA-seq data are shown in red, and were used to identify the TSS, which are indicated by curved black arrows on the annotation track. Annotated phage genes are grouped into functional clusters.

Transcription was observed from the promoter upstream of the *apI* gene (*STMMW_32112*) through to the uncharacterised gene *STMMW_32601* in certain conditions, including early stationary phase (ESP), bile shock and anaerobic shock. The corresponding genes in coliphage 186 belong to the early lytic operon (Shearwin and Egan, 2000) and represent the genes initially expressed during lytic phage replication. Transcription of a three-gene operon consisting of tail structural genes (*STMMW_31821, STMMW_31811* and *STMMW_31801*) was observed in a number of conditions and was particularly high in ESP, bile shock and nitric oxide shock. The tail structure of P2-like phages (Myoviruses) is complex (Christie and Calendar, 2016), and expression of these three genes alone would not produce functional phage tail particles. Therefore, the functional relevance of this transcript is unclear. Additionally, the *ogr* gene (*STMMW_31741*), reported to be involved in control of late gene expression in phage P2 (Christie and Calendar, 2016), was expressed in all conditions tested.

Despite the transcription of a subset of lytic genes in the BTP5 prophage, the repressor and integrase genes were transcribed in all conditions examined, albeit at low levels relative to the level of tail gene transcription. We conclude that the BTP5 prophage transcriptome does not inform the functionality of the prophage, and, consistent with our previous study, the lysogeny and lysis behaviour of the BTP5 prophage remains enigmatic (Owen et al., 2017). We speculate that there may be further control of prophage BTP5 gene expression at the post-transcriptional level, or alternatively the transcriptome may reflect heterogeneity in the activity of the BTP5 prophage across the bacterial cell population.

### Characterised prophage accessory loci have unique transcriptional signatures

*S.* Typhimurium prophages BTP1, Gifsy-2^D23580^, ST64B^D23580^ and Gifsy-1^D23580^ contain characterised accessory loci, including genes involved in *Salmonella* pathogenicity or phage exclusion (Figure 6A). To empirically determine whether known prophage accessory loci had distinct transcriptional signatures compared with the rest of the prophage, we used an expression cut-off of 100 TPM (Canals et al., 2019) to identify highly expressed genes. Genes with expression values of >100 TPM in at least one RNA-seq experiment were classified as highly expressed during lysogeny (Supplementary Table 4). The highly expressed prophage genes were assigned to one of the following functional categories based on annotation: unknown function, accessory gene, regulatory gene, integrase, transposase or structural gene (Figure 6B). Of the 278 genes annotated in the five prophages, 40 genes (14%) (Figure 6C) met our criteria of being highly expressed during lysogeny. As expected, a large number of the highly expressed category represented known accessory genes (11 genes), such as genes encoding type three secretion system (T3SS) effectors, or regulators (11 genes) including repressors. Among the other highly expressed genes were genes encoding two prophage integrases, one transposase, and one prophage structural protein. However, the largest category of highly expressed genes were of unknown function (14 genes) (Hinton, 1997). We conclude that the transcriptional signatures are consistent with these 14 genes being novel prophage accessory genes or regulatory genes. We note that this ‘guilt by association’ approach has previously been successfully used to identify novel *Salmonella* pathogenicity island (SPI)-regulated genes (Kröger et al., 2013), and to make broader regulatory deductions (Perez-Sepulveda and Hinton, 2018; Thattai Mukund, 2013).

**Figure 6.**
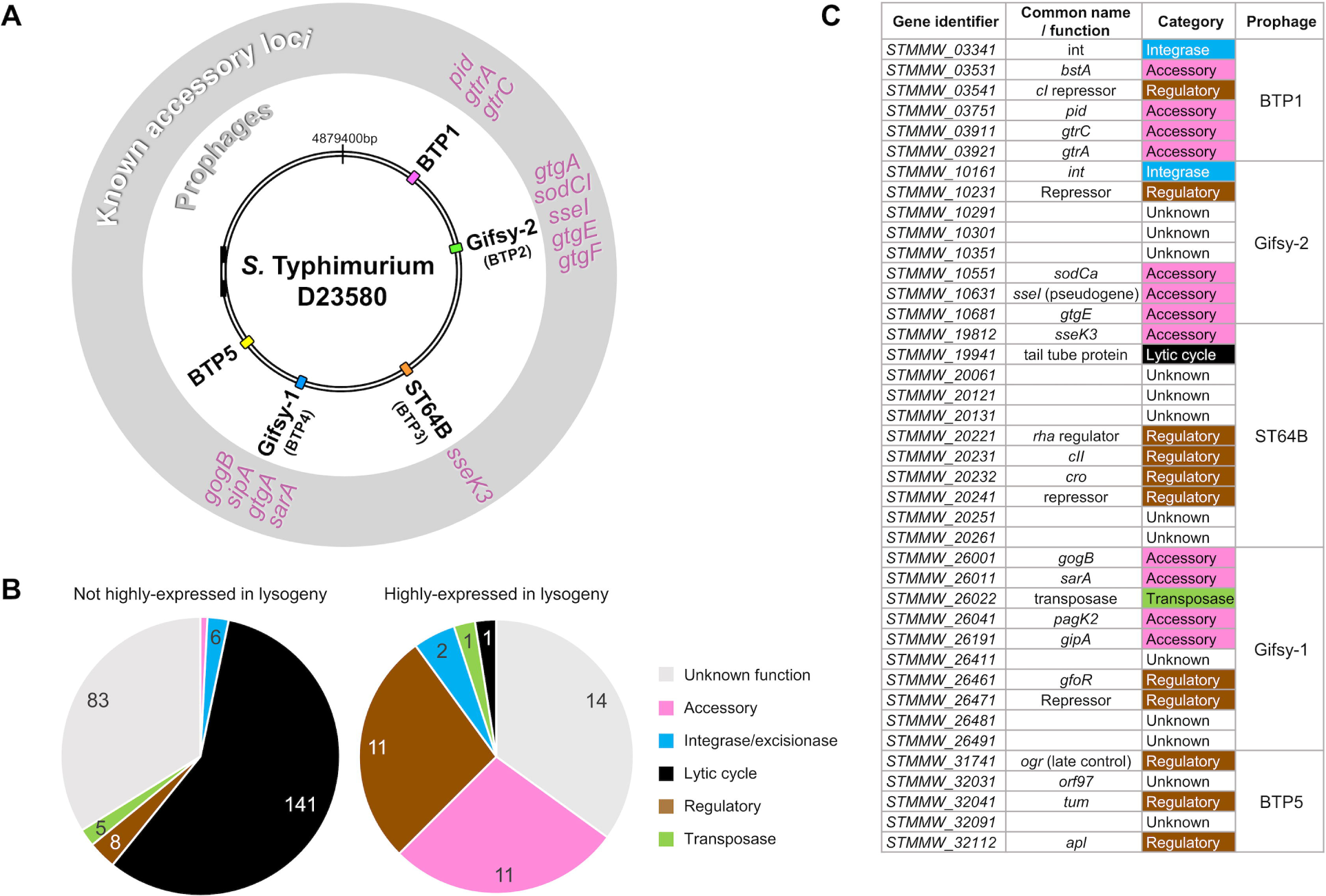
Prophage regulatory or accessory genes show unique transcriptional signatures. These findings suggest that transcriptomic data can be used to heuristically enrich for genes likely to be associated with novel regulatory or accessory functions. A. Genomic map of *S.* Typhimurium strain D23580 indicating the location of the five prophage elements. Known accessory loci associated with each prophage element are annotated in the exterior grey ring. B. Functional categorization of all prophage genes with expression values of <100 TPM (not highly expressed in lysogeny) and >100 TPM (highly expressed in lysogeny) in at least one RNA-seq condition. The majority of highly expressed prophage genes have known regulatory or accessory function, or have no known function. C. The 40 prophage genes of *S.* Typhimurium strain D23580 classified as highly expressed during lysogeny.

### Identification of putative novel prophage regulatory or accessory loci

Figure 7 shows the genomic and transcriptomic context of three of the prophage genes of unknown function identified in this study and likely to represent novel prophage regulatory or accessory loci. The *bstA* locus (Figure 7A) of prophage BTP1 is down-stream and co-transcribed with the *cI* repressor locus. The region between the repressor locus and the *n* locus of lambdoid prophages has been previously shown to have a high frequency of mosaicism and represents a common site of prophage accessory (moron) loci, such as the *rexAB* locus of phage lambda (Degnan et al., 2007). The *bstA* gene (*STMMW_03531,* formerly designated *ST313-td*) has been implicated as a determinant of both virulence (Herrero-Fresno et al., 2014) and anti-virulence (Herrero-Fresno et al., 2018) in *Salmonella enterica* strains, though no mechanism for these effects has been proposed (hence our conservative inclusion of *bstA* in the ‘unknown function’ functional category in this study. Our transcriptomic data support a functional role for *bstA* as a novel prophage accessory gene that deserves further study.

**Figure 7.**
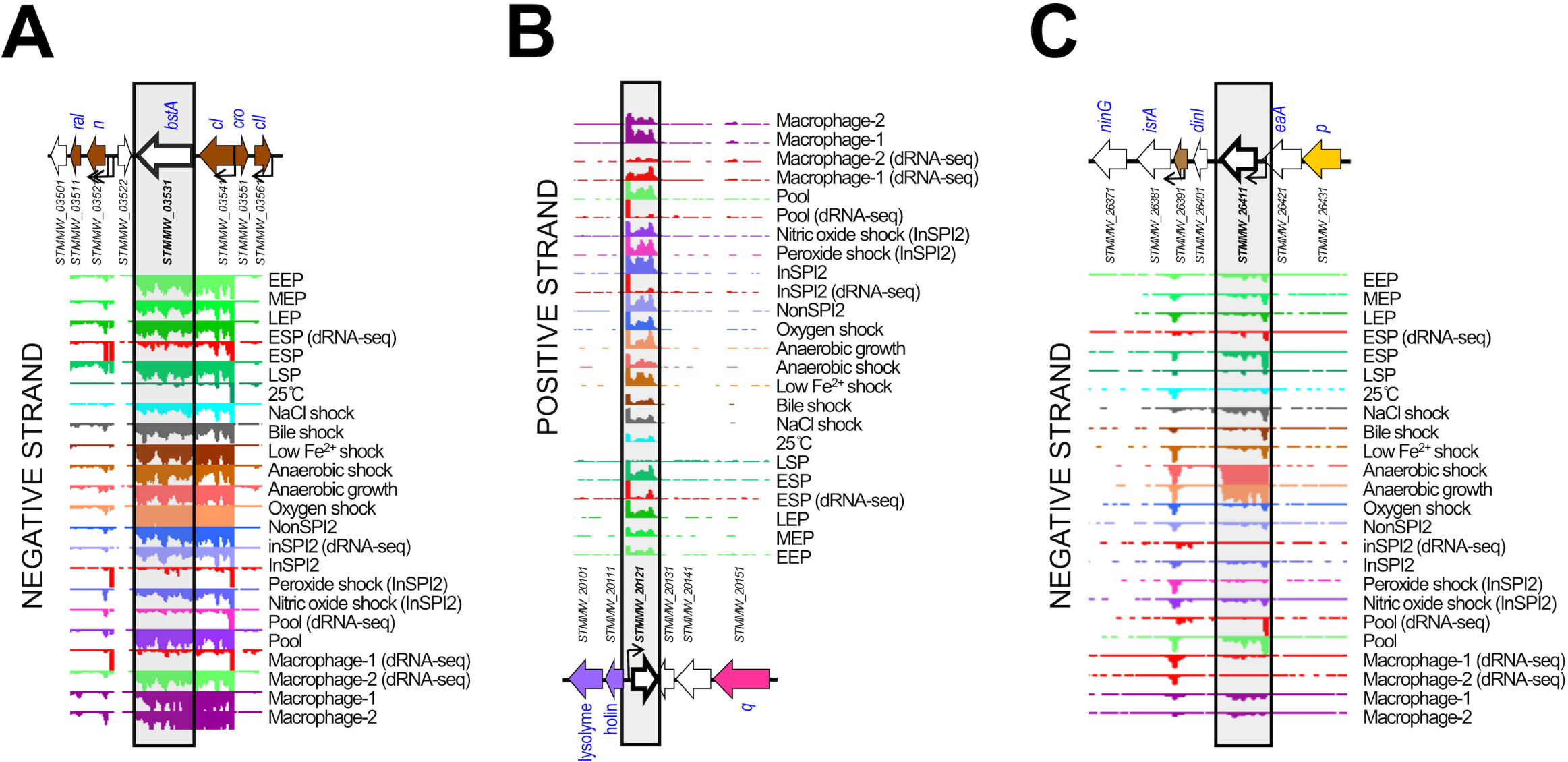
Examples of prophage genes exhibiting unique transcriptional signatures consistent with regulatory or accessory functions. A the *bstA* locus of prophage BTP1. B. The *STMMW_20121* locus of prophage ST64B. C. The *STMMW_26411* locus of prophage Gifsy-1,

Like *bstA*, the *STMMW_20121* locus (Figure 7B) of prophage ST64B^D23580^ showed transcription in almost all infection-relevant conditions included in our study. However, we observed that *STMMW_20121* was independently transcribed and terminated from its own promoter and terminator. *STMMW_20121* is encoded antisense to the prophage lytic genes, a region that is characteristic of prophage accessory (moron) loci (Cumby et al., 2012). The transcriptomic signature and genomic context of *STMMW_20121* is consistent with the gene product having an accessory or regulatory function.

Finally, we identified the *STMMW_26411* locus (Figure 7C) of prophage Gifsy-1^D23580^ as likely to be a novel prophage accessory gene. Unlike *bstA* and *STMMW_20121*, *STMMW_26411* showed highly condition-specific transcription, associated with anaerobic conditions. The environmental specificity of *STMMW_26411* transcription leads us to speculate that the gene is more likely to be a novel accessory gene than function as a prophage regulatory gene, particularly given the lack of corresponding lytic gene transcription in the Gifsy-1^D23580^ prophage under anaerobic conditions. This observation could be associated with the facultative anaerobic lifestyle of *S.* Typhimurium and the ability of the pathogen to colonise the mammalian gastrointestinal tract (Álvarez-Ordóñez et al., 2011). Furthermore, given the broad conservation of the Gifsy-1 prophage amongst *S.* Typhimurium strains (Mottawea et al., 2018) and known association of this prophage with virulence factors (Figure 6A), the *STMMW_26411* locus represents an exciting candidate accessory factor for further study.

### Identification of a novel prophage-encoded ncRNA involved in superinfection exclusion

As well as identifying novel candidate prophage accessory and regulatory genes, our transcriptomic analysis of the prophages of D23580 identified a number of putative novel ncRNAs. We focused on the ncRNAs of prophage BTP1 (Figure 1), as this prophage is functional (Owen et al., 2017), yet poorly characterised. To identify the biological relevance of novel ncRNAs, STnc6030 was selected for further investigation as it was particularly highly expressed in the majority of infection-relevant growth conditions (Figure 8A). Additionally, the putative ncRNA is unusually long, >700 nucleotides (nt) in length, and is positioned antisense to the BTP1 tailspike gene (*STMMW_03901*) and a gene encoding a putative DNA injection protein (*STMMW_03891*). It should be noted that the STnc6030 region is an area of unusually high transcription, with 10 TSS (sense and antisense) defined within the tailspike gene alone. Due to the length and antisense location of the STnc6030 transcript, the putative ncRNA was hypothesised to be an asRNA species. The majority of asRNAs that have been identified in bacteria function as inhibitors of target RNA function (Wagner et al., 2002), and are commonly found in accessory genome elements such as phages and plasmids (Thomason and Storz, 2010). We hypothesised that STnc6030 interacts with the transcription of the BTP1 tailspike gene, and investigated this experimentally. Figure 8A shows a detailed view of STnc6030 transcription within the BTP1 transcriptome. Analysis of the STnc6030 transcript region in all three reading frames did not reveal any open reading frames >60 amino acids in length, supporting the classification of STnc6030 as an ncRNA species. The beginning of the STnc6030 transcript corresponds to the beginning of RNA-seq reads mapping to the tailspike gene on the sense strand, consistent with antisense interference with the tailspike gene transcript.

**Figure 8.**
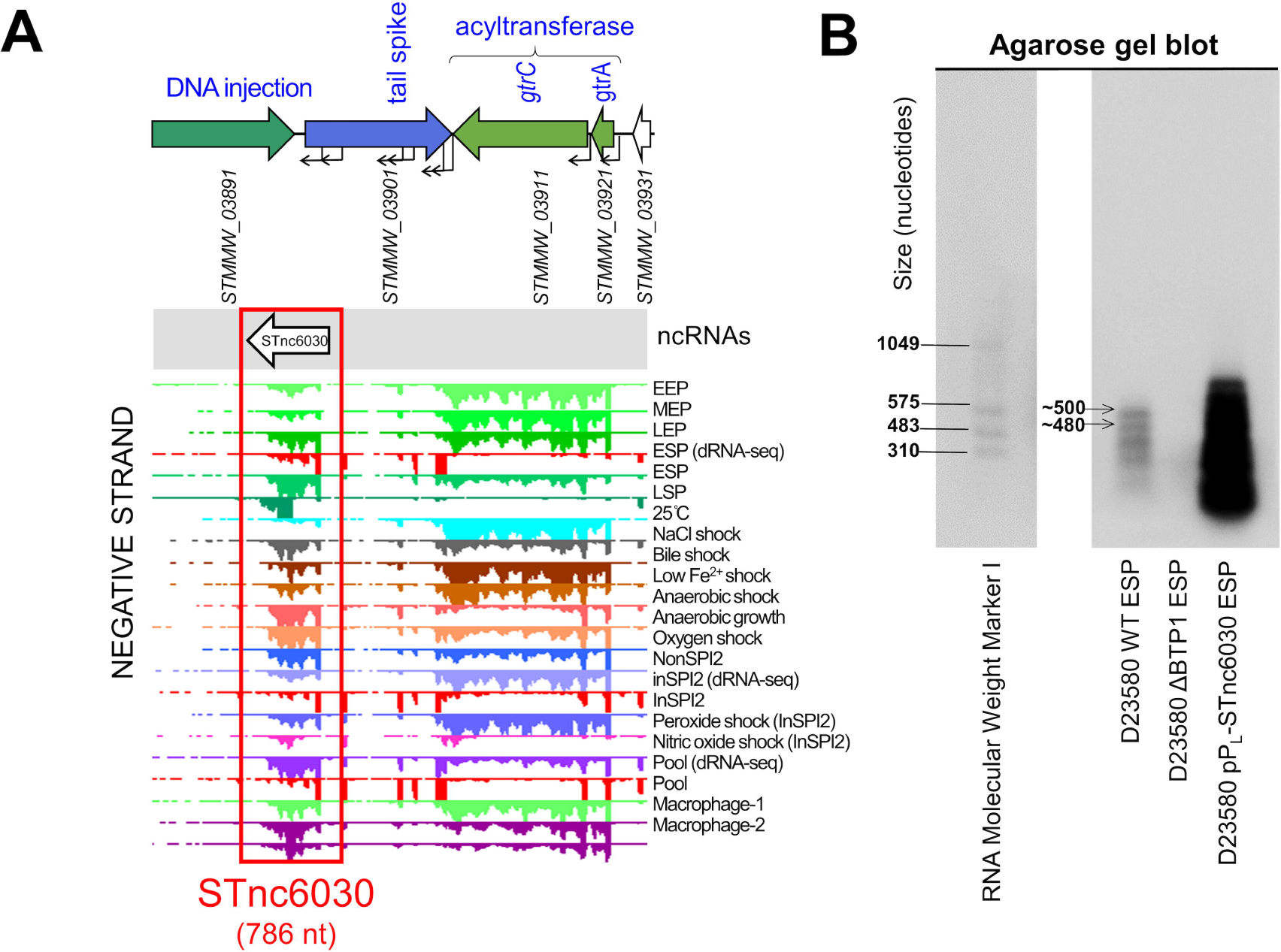
Transcriptomics-based identification of a novel prophage-encoded asRNA, STnc6030. A. The transcriptomic context of the STnc6030 ncRNA. The transcriptomic data are the same as shown in Figure 1. B. Detection of the STnc6030 transcript by northern blot using an anti-STnc6030 DIG-labelled riboprobe. The two most abundant transcripts detected by the anti-STnc6030 riboprobe are indicated, with the approximate size estimated from the molecular weight ladder.

The putative STnc6030 transcript that was identified from the BTP1 transcriptomic data was cloned into the pP_L_ expression plasmid under the control of the constitutive P_LlacO-1_ promoter. To confirm the presence of the STnc6030 transcript, an anti-STnc6030 riboprobe was synthesised to detect the STnc6030 RNA species by Northern blot (Figure 8B). The riboprobe was designed to cover the totality of the approximately 786 nt STnc6030 transcript, allowing the detection of any transcripts corresponding to this region. The anti-STnc6030 riboprobe detected a number of transcripts in the D23580 wild-type (WT) strain, whilst no transcripts were detected in the D23580 ΔBTP1 mutant, confirming that the detected bands did not reflect non-specific binding. The largest transcript detected by the anti-STnc6030 probe in the D23580 WT was approximately 500 nt in length, significantly shorter than the putative length identified from the transcriptomic data (786 nt). At least two other smaller transcripts were detected, of ∼500 nt and ∼480 nt in length which could result from RNA processing or degradation product, as seen for other *Salmonella* ncRNAs such as ArcZ (Papenfort et al., 2009).

To interrogate the biological function of STnc6030, D23580 WT and D23580 ΔBTP1 (sensitive to phage BTP1) were transformed with the pP_L_-STnc6030 and empty vector control plasmids. The BTP1 prophage displays an unusually high level of spontaneous induction in the lysogeny cell population (Owen et al., 2017) and we hypothesised that STnc6030 may contribute to this phenotype. However, over-expression of STnc6030 RNA did not affect the level of BTP1 spontaneous induction in the D23580 WT background, with no difference in the number of spontaneously induced BTP1 phage in overnight culture supernatants of D23580 WT, D23580 pP_L_-STnc6030 and D23580 pP_L_ (empty vector) (Figure 9A).

**Figure 9.**
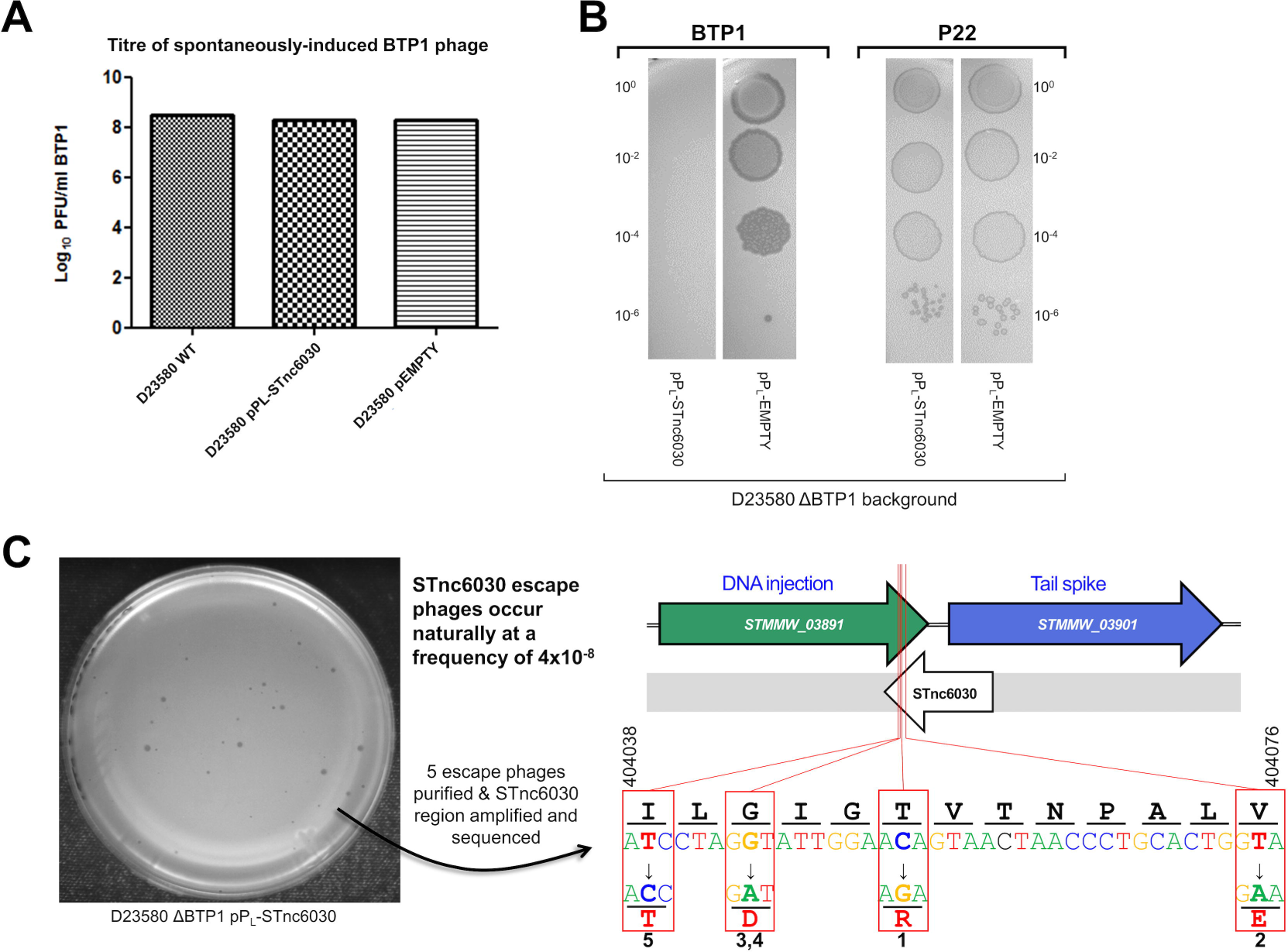
The BTP1-prophage encoded asRNA STnc6030 functions as a phage-specific super-infection exclusion factor. A. Over-expression of the STnc6030 RNA in D23580 WT background does not affect BTP1 spontaneous induction. Plaque assay of overnight culture supernatants of D23580 WT, D23580 pP_L_-STnc6030 and D23580 pP_L_ (empty vector) on host strain D23580 ΔBTP1. B. Heterologous expression of STnc6030 in D23580 ΔBTP1 completely protects against BTP1 phage, but not P22 infection. Plaque assay of BTP1 and P22 phage on D23580 ΔBTP1 strains containing the pP_L_-STnc6030 expression plasmid or the negative control plasmid. C. Isolation of STnc6030-escape mutants of phage BTP1 Suggest the STnc6030 functional ‘seed’ region is located at the 3’ end of the transcript. A high titer BTP1 phage stock was used to identify naturally occurring BTP1 phage mutants that were immune to inhibition by STnc6030. Escape phages were estimated to occur at a frequency of approximately 4x10^-8^. Five escape phages of varying plaque morphologies were selected for sequencing. The sequence of the STnc6030 region of the escape phages identified SNPs. The position and substitution are shown. The SNPs identified that conferred immunity to STnc603*0* interference were clustered within a 36bp region (shown) corresponding to the 3’ end of the STnc6030 transcript and the 3’ end of the *STMMW_03891* gene (located between nt 404040 and 404076 on the D23580 chromosome Accession: FN424405).

Next, we investigated whether STnc6030 could play a role in phage immunity. Expression of STnc6030 in a naïve host in the absence of the BTP1 prophage (D23580 ΔBTP1 pP_L_-STnc6030) mediated total immunity to BTP1 infection (Figure 9B), but did not modulate susceptibility to phage P22 infection. These results were consistent with a regulatory mechanism in which STnc6030 targeted the sense transcript of the BTP1 DNA injection and tailspike genes for degradation by base pairing. RNA-RNA interactions require high nucleotide complementarity, and consistent with this model, the corresponding region of the P22 genome does not share similarity to BTP1 at the nucleotide level.

To explore the functionality of the STnc6030 transcript, a high titre BTP1 phage stock was screened for the presence of naturally occurring mutants resistant to exclusion by STnc6030 (Figure 9C). The BTP1 phage stock was plated on lawns of D23580 ΔBTP1 containing the pP_L_-STnc6030 expression plasmid escape phages arose at a frequency of 4×10^-8^, and had varying plaque sizes (Figure 9C). Five escape phages were isolated and purified, and a nested PCR strategy was used to amplify the STnc6030 transcript region of the escape phages. Sequencing of the STnc6030 region revealed that each escape phage contained a single nucleotide polymorphism (SNP) relative to the BTP1 WT sequence (Figure 9C). A total of four unique SNPs were identified that conferred resistance to STnc6030-mediated exclusion. The four SNPs clustered within a 36 bp region corresponding to the 3’ end of the STnc6030 transcript, antisense to the putative DNA injection gene *STMMW_03891* (Figure 9C). These data implicate the 3’ end of STnc6030 as a functionally active “seed” region of the RNA which interacts with the antisense target (*STMMW_03891*). The 4 SNPs also cause non-synonymous amino acid substitutions to the STMMW_03891 protein (Figure 9C).

Together, these results are consistent with a model where the STnc6030 asRNA acts as a superinfection exclusion factor, inhibiting re-infection of the BTP1 phage lysogen with ‘self’ phage, or preventing the infection of closely-related phages that share sequence identify to BTP1 in the STnc6030 region. However, the fact that the BTP1 prophage induction can occur normally in the presence of the STnc6030 asRNA (Figure 9A) suggests that the induced BTP1 prophage has a mechanism to escape the effects of its own superinfection exclusion RNA. Further work is required to confirm the biological activity of the STnc6030 asRNA of the BTP1 prophage. Our data illustrate the power of using transcriptomic data to uncover novel prophage biology. Indeed, ncRNA species are completely undetectable without combining prophage genome sequence data with corresponding transcriptomic data of the lysogen.

## DISCUSSION

Prophage accessory loci are often responsible for lysogenic conversion of the bacterial host, such as genes encoding virulence factors or superinfection immunity factors (Casjens and Hendrix, 2015). A study in *Pseudomonas aeruginosa* showed that 12 out of 14 previously uncharacterised accessory (moron) loci affected diverse bacterial phenotypes including phage immunity, motility and biofilm formation (Tsao et al., 2018). However, prophage accessory genes are difficult to identify from DNA-sequence alone. Prophage accessory loci are likely to be associated with unique transcriptional signatures compared to phage lytic genes, because in order to affect the biology of the host cell, they must be expressed during lysogeny. Here, we show that transcriptomic data, pre-existing or purposefully generated, provide unique insights into the molecular biology of prophages, and we propose that transcriptional signatures should be used to reduce the challenge of identifying novel prophage regulatory and accessory genes.

Inferences from the transcriptomes of the D23580 prophage regions were generally consistent with our previous findings concerning the functionality of the prophages (Owen et al., 2017). BTP1, a prophage which exhibits an unusually high level of spontaneous induction, showed an unusually high level of transcriptional activity for a prophage region. Canonically, prophage genes are expected to be transcriptionally repressed during lysogeny, apart from those genes whose products maintain prophage lysogeny, such as the gene encoding the CI repressor in lambdoid prophages (Ptashne, 2004) or genes with accessory functions that confer a fitness advantage to the lysogen. Unlike the four other prophage regions of D23580, low-level transcription of the BTP1 structural genes was observed in almost all growth conditions tested. A lysogenic cell cannot constitutively express phage structural and lytic genes, because, once the prophage molecular switch has moved to lytic from lysogenic replication, the unavoidable consequence is cell death (Ptashne, 2004). Therefore, if the observed lytic gene expression occurred in the entire cellular population, a population collapse would ensue, as ultimately sufficient lysis proteins were accumulated to initiate cell lysis. However, the BTP1 prophage lysogen (D23580) exhibits normal growth dynamics that are comparable to strains not lysogenised by the BTP1 prophage (Owen et al., 2017). We therefore speculate that lytic gene expression observed in the transcriptomic data reflects the unavoidable averaging of gene expression across a heterogeneous population which has very high lytic gene expression in the subset of the bacteria undergoing spontaneous BTP1 prophage induction.

Consistent with this speculation, the remaining D23580 prophages (Gifsy-2^D23580^, ST64B^D23580^, Gifsy-1^D23580^, and BTP5) do not exhibit significant spontaneous induction levels (Owen et al., 2017) and show very little lytic gene expression in the majority of growth conditions (Figures 2, 3, 4 & 5). The only other D23580 prophage to show evidence of late gene transcription was ST64B^D23580^ in two growth conditions (peroxide and nitric oxide stress) and could reflect specific induction behaviour of the ST64B^D23580^ prophage. In light of this finding, we speculate that the absolute expression levels of prophage structural genes could be used *in silico* to quantify the fraction of the lysogenic population undergoing lytic prophage replication.

As well as providing insight into the replication state of the D23580 prophages, the transcriptome maps also allow the identification of putative accessory regions, genes or transcripts, expressed during lysogenic replication, or represent novel genes involved in the regulation of lysogeny. Prophage accessory genes are of importance for bacterial pathogens, as they could represent ‘smoking guns’ that could be responsible for rapid changes in disease tropism. Prophages BTP1 and BTP5 are specific to the epidemic African sequence type of *S.* Typhimurium ST313, and therefore have not been well characterised. The transcriptome maps of prophages BTP1 and BTP5 showed several more regions of transcription than are theoretically necessary for a lambdoid prophage to maintain lysogeny (typically the *cI* repressor only) (Hendrix, 1983).

A number of genes in the BTP1 prophage showed an expression pattern consistent with an accessory or regulatory function, including *bstA*, *pid*, *gtrA^BTP1^* and *gtrC^BTP1^*. Of these four genes, all except *bstA* have mechanistically described accessory functions. Prophages Gifsy-2, ST64B and Gifsy-1 are broadly conserved in many strains of *S. enterica* (Mottawea et al., 2018) and encode numerous virulence genes including T3SS effector genes (Ehrbar and Hardt, 2005). Although the regulatory behaviour and accessory genes of these prophages have been well studied, we identified numerous novel candidate accessory and regulatory genes, demonstrating the power of our transcriptomic approach.

Whilst transcriptomics represents a useful tool for discovering coding gene functions, it is arguably an even more powerful approach for the discovery of non-coding genomic elements such as ncRNAs. A number of putative RNA transcripts in the BTP1 prophage which did not correspond to protein coding sequences had an expression pattern consistent with an accessory function, including eight putative novel ncRNAs (Canals et al., 2019). Several prophage-encoded ncRNAs have been implicated in bacterial virulence, for example, the Gifsy-1 prophage encoded IsrJ, a ∼74 nt ncRNA required for efficient invasion of *Salmonella* into nonphagocytic cells and effector translocation by the SPI-1 T3SS (Padalon-Brauch et al., 2008). Prophage-encoded ncRNAs also mediate non-virulence accessory functions, including the *sas* asRNA of phage P22 that induces a translational switch between distinct peptides encoded by the *sieB* gene, and is critical to the function of the SieB superinfection exclusion system (Ranade and Poteete, 1993). Lastly, the phage λ ncRNA OOP inhibits CII protein synthesis thereby pushing the phage molecular decision towards lysis, rather than lysogeny (Krinke and Wulff, 1987).

Our transcriptomic approach identified the STnc6030 asRNA encoded within the BTP1 prophage late genes. The transcript is located antisense to the 3’ end of the putative DNA-injection gene *STMMW_03891* and 5’ end of the tailspike gene *STMMW_03901*. Expression of STnc6030 in a heterologous host abolished susceptibility to infection by BTP1, but not the related phage P22. However, the spontaneously induced titre of BTP1 phage was not affected when the STnc6030 transcript was overexpressed, suggesting that the asRNA does not interfere with replication of spontaneously induced BTP1.

Bacterial asRNAs, also known as *cis*-encoded RNAs, are usually found antisense to annotated coding genes. The extensive genetic complementarity with the corresponding transcripts allows asRNAs to affect the stability of complementary mRNA transcripts by base-pair interactions (Gottesman and Storz, 2011). Because double-stranded RNA molecules are substrates for endoribonucleases, the effect of asRNA targeting is usually to increase the degradation of particular mRNA transcripts and so reduce levels of the gene product. Alternatively, base-pairing of two RNA species can block an RNase recognition site, leading to increased stability of the target mRNA (Thomason and Storz, 2010). Our data are consistent with a model where the functional mechanism of the STnc6030 asRNA is a base-pairing interaction with the transcript containing the DNA-injection gene *STMMW_03891*, decreasing the stability of the mRNA. As prophage genes are frequently expressed as polycistronic operons encoding many of the genes required by the replicating phage, the antisense targeting of a single prophage gene could destabilize a long transcript encoding the entirety of the prophage lysis and structural genes, effectively inhibiting prophage replication. However, it remains unclear how the asRNA, which is natively located within BTP1, avoids interference with the BTP1 prophage upon induction from lysogenic growth within the bacterial chromosome. The mechanism by which the induced BTP1 prophage escapes its own immunity asRNA remains to be discovered. The BTP1 prophage encodes two other systems for superinfection exclusion, the GtrAC^BTP1^ system and the CI^BTP1^ repressor (Kintz et al., 2015), so the precise biological role of the STnc6030 asRNA in the context of these other systems requires further study.

Overall, our work represents the first detailed report of the transcriptional landscapes of native bacterial prophages. A wealth of RNA-sequencing data exist for a number of poorly characterised bacterial pathogens in which virulence factors and prophage molecular regulation have not been characterised. As well as identifying novel candidate regulatory and accessory loci in *Salmonella* prophages, our work represents a ‘proof of concept’ study that shows that careful analysis of RNA-seq data mapped to prophage regions could reveal a vast array of novel putative prophage accessory loci. Prophage transcriptomic maps represent a powerful window through which to view the molecular biology of temperate phages.

## AUTHORSHIP & CONTRIBUTIONS

SVO and JCDH conceptualised the study. SVO, RC, DH and CK developed methodology, curated data, and contributed resources. SVO, RC and NW conducted investigation and formal analysis. RC, NW, DH, CK and JCDH provided supervision. SVO wrote the manuscript. All authors reviewed and edited the manuscript.

## ACKNOWLEDGEMENTS

We are grateful to present and former members of the Hinton laboratory and Heather Allison for helpful discussions. We thank Paul Loughnane for expert technical assistance, and Michael Baym for his patience.

## Figure Legends

Supplementary Figure 1. Heat map showing the absolute expression of BTP1 genes in 17 infection-relevant conditions. RNA-seq data from Canals et al., 2019 were used to generate absolute expression values (transcript per million, TPM) for each coding gene and annotated ncRNA of the BTP1 prophage.

Supplementary Figure 2. Heat map showing the absolute expression of the Gifsy-2 prophage of D23580 and 4/74 strains in 17 infection-relevant conditions. RNA-seq data from Canals et al., 2019 were used to generate absolute expression values (transcript per million, TPM) for each coding gene and annotated ncRNA of the Gifsy-2 prophage. Ortholog IDs in red text indicate genes unique to Gifsy-2^D23580^ or Gifsy-2^4/74^.

Supplementary Figure 3. Heat map showing the absolute expression of the ST64B prophage of D23580 and 4/74 strains in 17 infection-relevant conditions. RNA-seq data from Canals et al., 2019 were used to generate absolute expression values (transcript per million, TPM) for each coding gene and annotated ncRNA of the ST64B prophage. Ortholog IDs in red text indicate genes unique to ST64B^D23580^ or ST64B^4/74^.

Supplementary Figure 4. Heat map showing the absolute expression of the Gifsy-1 prophage of D23580 and 4/74 strains in 17 infection-relevant conditions. RNA-seq data from Canals et al., 2019 were used to generate absolute expression values (transcript per million, TPM) for each coding gene and annotated ncRNA of the Gifsy-1 prophage. Ortholog IDs in red text indicate genes unique to Gifsy-1^D23580^ or Gifsy-1^4/74^.

Supplementary Figure 5. Heat map showing the absolute expression of BTP5 genes in 17 infection-relevant conditions. RNA-seq data from Canals et al., 2019 were used to generate absolute expression values (transcript per million, TPM) for each coding gene of the BTP5 prophage.

