## Supplementary figures and images for "A window into lysogeny: Revealing temperate phage biology with transcriptomics"

### Supplementary Figure 1

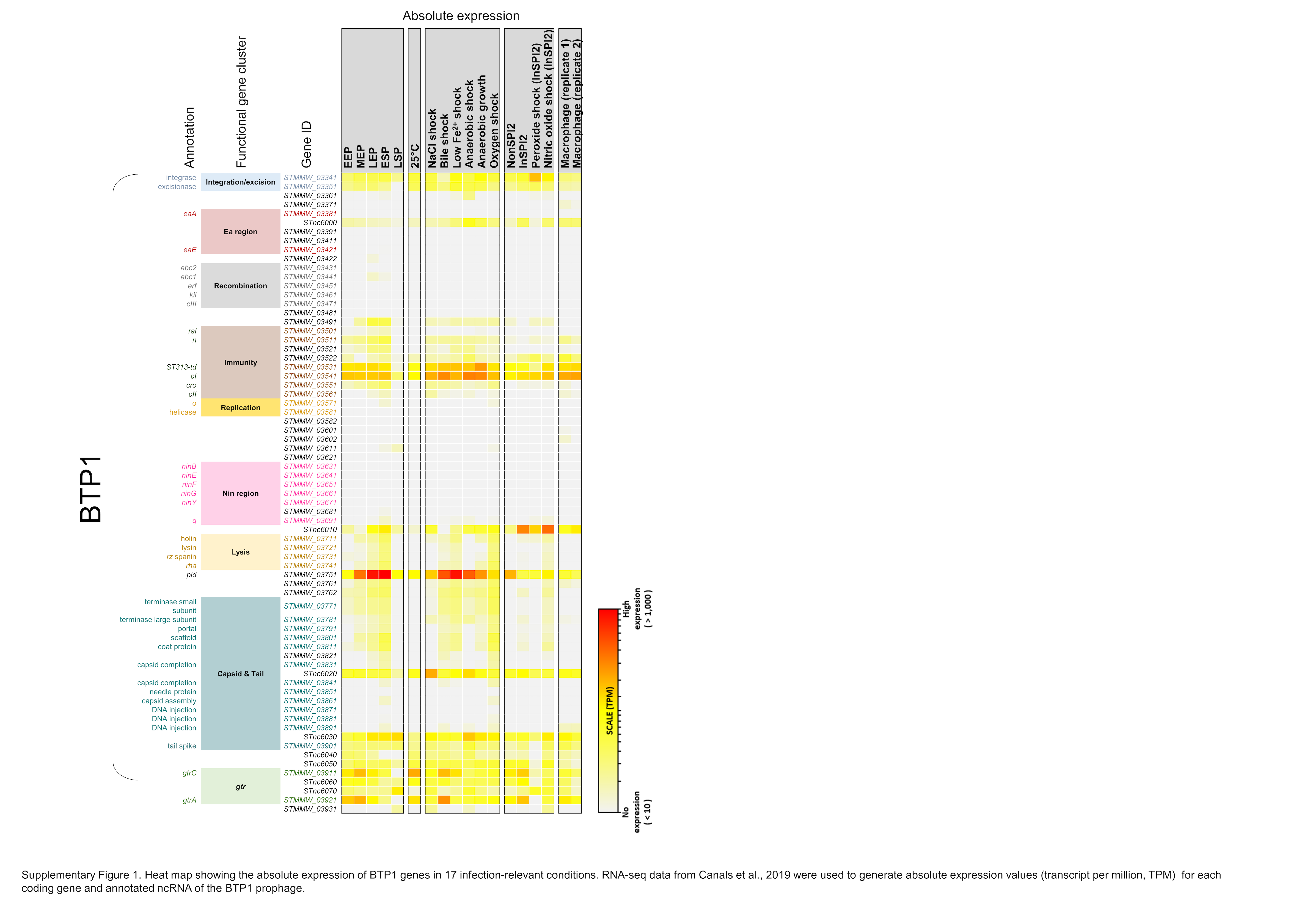

### Supplementary Figure 2

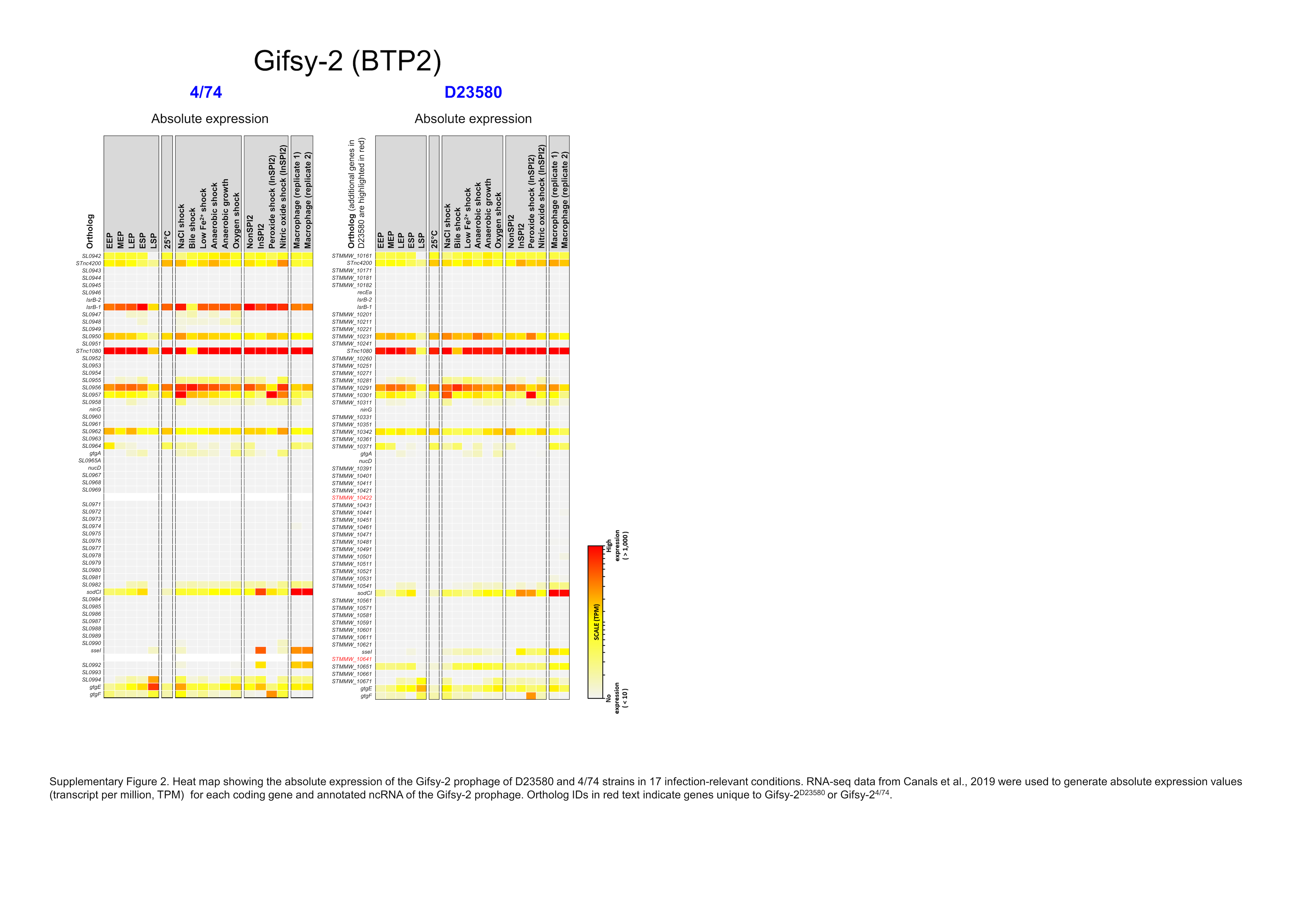

### Supplementary Figure 3

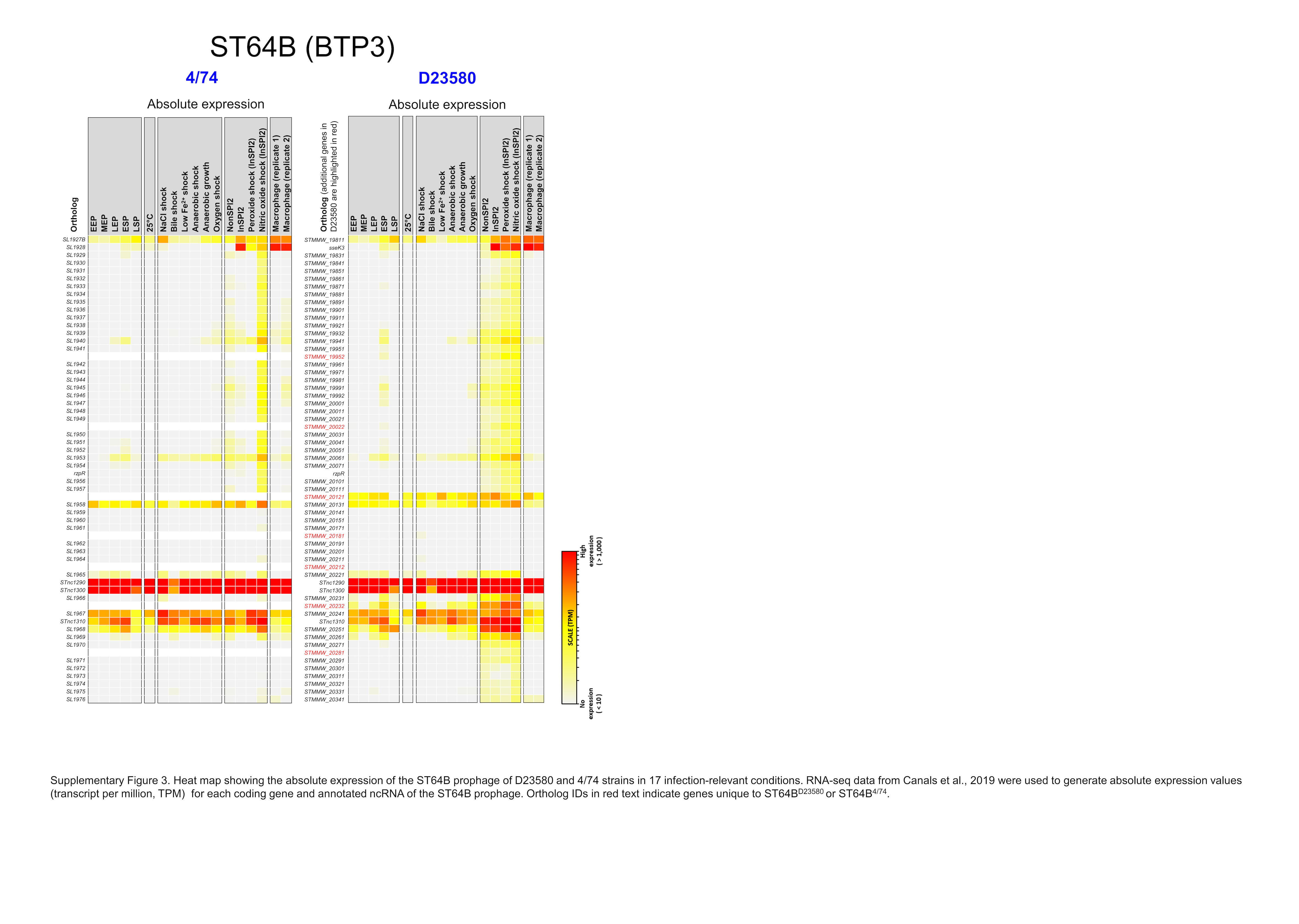

### Supplementary Figure 4

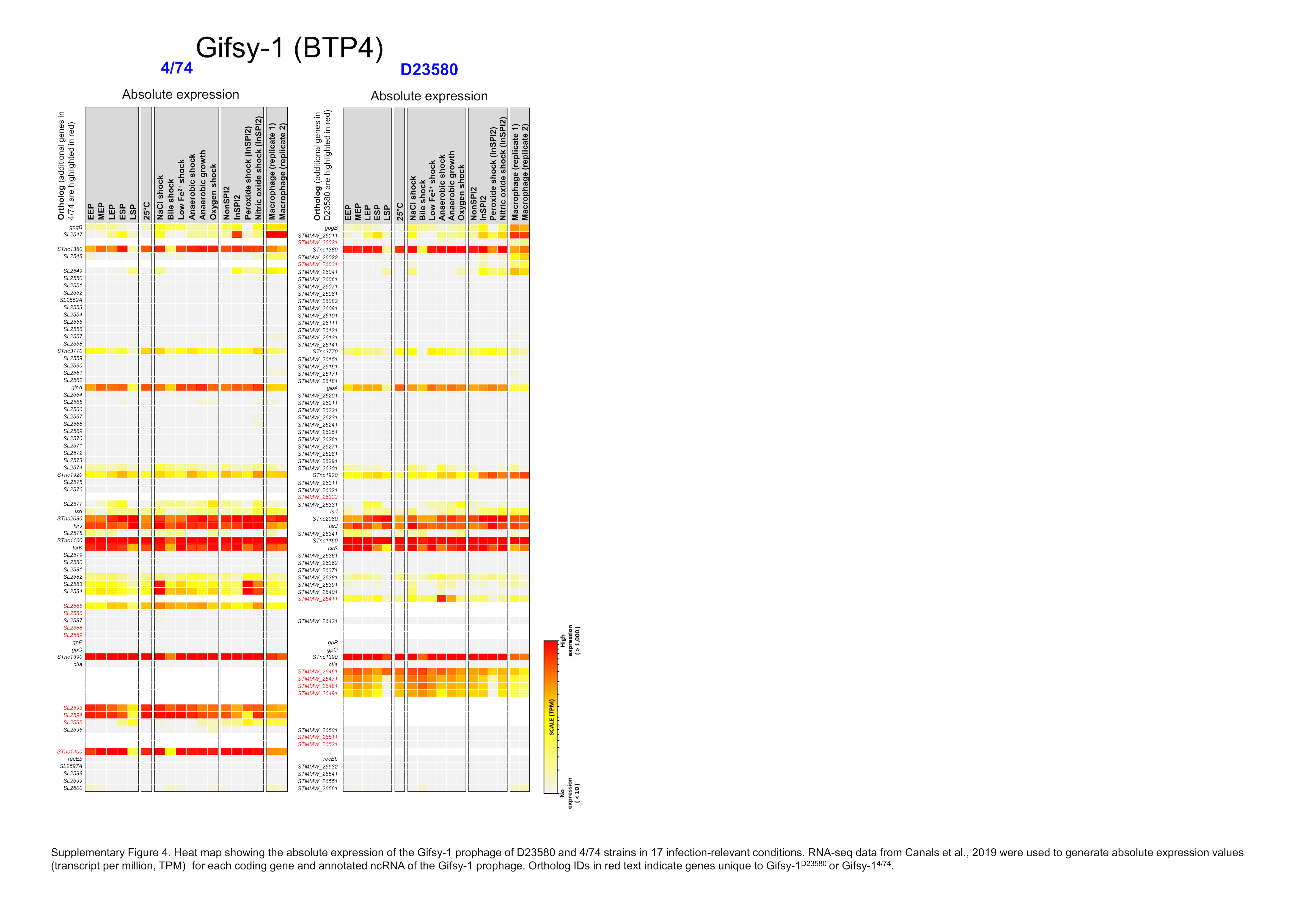

### Supplementary Figure 5

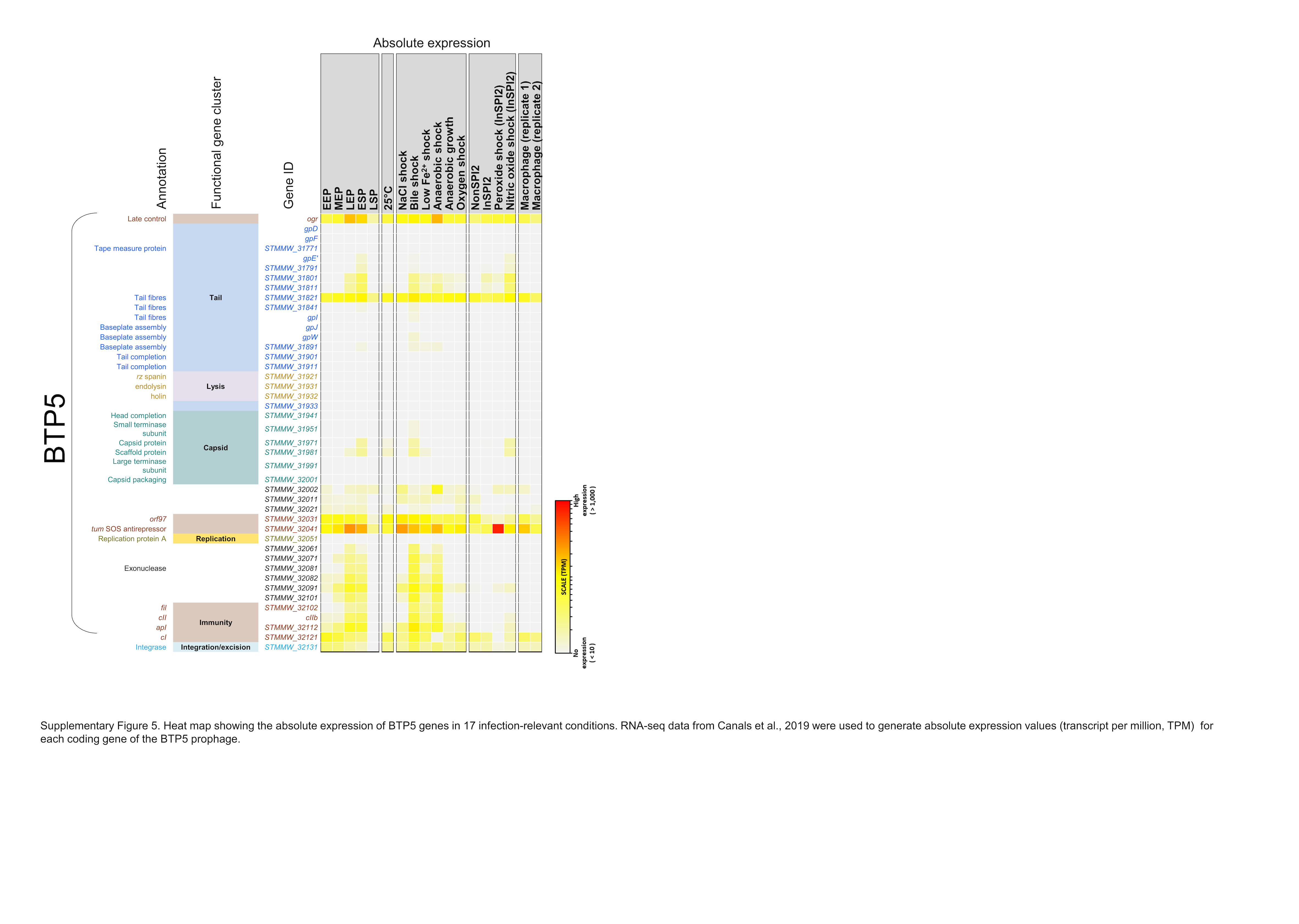
