## Supplementary Tables 1-3 for "A window into lysogeny: Revealing temperate phage biology with transcriptomics"

| **RNA-seq experiment** | **Abbreviation** | **Growth conditions** |
| --- | --- | --- |
| Early exponential phase | EEP | Growth in Lennox broth to OD_600_ 0.1 |
| Mid exponential phase | MEP | Growth in Lennox broth to OD_600_ 0.3 |
| Late exponential phase | LEP | Growth in Lennox broth to OD_600_ 1.0 |
| Early stationary phase | ESP | Growth in Lennox broth to OD_600_ 2.0 |
| Late stationary phase | LSP | Growth in Lennox broth to OD_600_ 2.0 + 6 h |
| Low temperature | 25°C | Growth in Lennox broth to OD_600_ 0.3 at 25°C (It's growth at low temperature (ON culture 37°C, 200 RPM, Lennox, diluted 1:1000 and grown at 25 °C until OD_600_ = 0.3) |
| Osmotic shock | NaCl_S | Growth in Lennox broth to OD_600_ 0.3; then addition of NaCl to a final concentration of 0.3 M for 10 min |
| Addition of bile | Bile_S | Growth in Lennox broth to OD_600_ 0.3; then addition of bile to a final concentration of 3% for 10 min |
| Iron limitation | LwFe_S | Growth in Lennox broth to OD_600_ 0.3; then addition of 2,2’-dipyridyl to a final concentration of 0.2 mM for 10 min |
| Anaerobic shock | No_O2_S | Growth in Lennox broth to OD_600_ 0.3 (50 ml), then filled into 50 ml closed centrifuge tube and incubated without agitation for 30 min at 37°C (Falcon tube) |
| Anaerobic growth | No_O2 | Static growth in Lennox broth to OD_600_ 0.3 in a completely filled and closed 50 ml centrifuge tube (Falcon tube) |
| Aerobic shock | O2_S | Static growth in Lennox broth to OD_600_ 0.3 in a completely filled and closed 50 ml centrifuge tube (Falcon tube); then 15 min aerobic growth (baffled flask, 250 rpm) |
| SPI2 non-inducing conditions | noSPI2 | Growth in PCN medium to OD_600_ 0.3 (pH 7.4, 25 mM Pi) |
| SPI2 inducing conditions | inSPI2 | Growth in PCN medium to OD_600_ 0.3 (pH 5.8, 0.4 mM Pi) |
| Oxidative stress | H2O2_S | PCN to OD_600_ 0.3, then addition of H_2_O_2_ to final concentration of 1 mM H_2_O_2_ for 12 min |
| Nitric oxide | NOs | Growth in PCN medium to OD_600_ 0.3 (pH 5.8, 0.4 mM Pi); then addition of 250 μM Spermine NONOate for 20 min |
| RNA from macrophages | MAC | RNA isolated from RAW 264.7 murine macrophages 8 h post infection |

Supplementary Table 1. Description of RNA-seq experiment growth conditions as described in Kröger et al., 2013 and Canals et al., 2019.

| Oligo Name | Sequence (5’🡪3’) | Purpose |
| --- | --- | --- |
| NW_88 | CTAAATACATTCAAATATGTATCCGGTCCAACCAGCGGCACCAG | Replacement of *bla* (ap^R^) by *aacC1* (Gm^R^ ) in pP_L_ |
| NW_89 | GTAAACTTGGTCTGACAGTTACCAATTAGGTGGCGGTACTTGGG | Replacement of *bla* (ap^R^) by *aacC1* (Gm^R^ ) in pP_L_ |
| STnc6030_pPL_F (NW_295) | GTGAGCGGATAACAAGATACTGAGCACAGCAATATAGTCAACCTGAGAAC | For insertion of STnc6030 asRNA into expression plasmid pPL-Gm using overlap extension PCR cloning |
| STnc6030_pPL_R (NW_296) | GCCTTTCGTTTTATTTGATGCCTCTAGACTGCGTATCTGAAGGGGATTAAG |  |
| STnc6030_T7 (NW_297) | TAATACGACTCACTATAGGGCTGCGTATCTGAAGGGGATTAAG | For synthesis of anti-STnc6030 riboprobe. Use with STnc60_pPLGM_f |
| STnc6030_pPL_R NW_296 | GCCTTTCGTTTTATTTGATGCCTCTAGACTGCGTATCTGAAGGGGATTAAG | Amplification of STnc6030 region from BTP1 prophage |
| NW_298 | TAATACGACTCACTATAGGGCATAGTCAGGAAGAAGTGAT |  |
| STnc6030_pPL_F  NW_295 | GTGAGCGGATAACAAGATACTGAGCACAGCAATATAGTCAACCTGAGAAC | Used with NW_296 for nested amplificiation of STnc6030 region from amplification product of NW_298 and NW_295 |

Supplementary Table 2. All oligonucleotide sequences used in this study

Supplementary Table 3. All bacterial strains, phages and plasmids used in this study

| **Type** | **Name** | **Description** | **Reference / origin / accession (if available)** |
| --- | --- | --- | --- |
| Bacterial strain | JH3621 | *S.* Typhimurium D23580 WT | (Kingsley et al., 2009) / FN424405 |
| Bacterial strain | JH3810 | *S.* Typhimurium D23580 ΔBTP1 | (Owen et al., 2017) |
| Bacterial strain | JH3676 | *S.* Typhimurium 4/74 WT | (Rankin and Taylor, 1966) / CP002487 |
| Bacterial strain | *E. coli* TOP10 | Highly competent *E. coli* strain for cloning procedures | Invitrogen |
| Phage | BTP1 WT |  | Isolated from D23580 supernatant / LT714109.2 |
|  | P22 WT |  | Gift from A. Aertsen / BK000582 |
| Plasmid | pP_L_ (pJV300) | Expression plasmid, Ap^R^ | (Sittka et al., 2007) |
| Plasmid | pME4510 | Promoter search vector pME4510, Gm^R^ | (Rist et al., 1998) |
| Plasmid | pP_L_-STnc6030 | Expression plasmid; STnc6030 sRNA under the control of the P_LlacO-1_ promoter; Gm^R^ | This study |
| Plasmid | pP_L_-Gm | Derived from pJV-300 but *bla* gene replaced with *aacC1* gene from pME4510; Gm^R^ | This study |
